# Evidence of positive polygenic selection following a series of environmental and demographic disasters in Pacific herring

**DOI:** 10.1101/2025.08.05.668748

**Authors:** Kaho H. Tisthammer, Joseph A. McGirr, Elias M. Oziolor, Vince Buffalo, Jennifer Roach, Andrew Whitehead

## Abstract

The causes and consequences of population collapse and recovery are difficult to discern and therefore predict because historical data are often sparse, and environmental drivers may interact in complex ways. Yet the influence of environmental change can imprint in contemporary and historic DNA sequences, thereby offering clues about drivers of population dynamics. The Prince William Sound (PWS) stock of Pacific herring famously collapsed in the early 1990s following the *Exxon Valdez* oil spill (EVOS), yet the causes of the collapse and reasons for a persistent lack of recovery are controversial. Three seasons after the spill, the PWS population declined precipitously, coincident with repeated disease outbreaks, and has remained small across the three decades since. We use population genetics as a forensic tool to gain insight into the causes and consequences of the collapse. We tracked genome-wide genetic variation across time, spanning the period immediately after the spill but preceding the collapse (1991) and across three subsequent decades to 2017. We compared genetic change through time between the PWS population and two reference Alaska populations. We detected high genetic differentiation between the Bering Sea and the Gulf of Alaska, and relatively small but significant genetic structure within the Gulf of Alaska. No loss of genome-wide genetic diversity was observed in the PWS population after the collapse. We did not detect any evidence for selective sweeps at individual loci. However, using sensitive temporal covariance methods, we detected significant signatures of positive polygenic adaptation in the PWS population but not in either of the reference populations. Gene functional enrichment analysis is consistent with selection acting on immune function and pathways that regulate crude oil developmental toxicity in the period following the spill and collapse. This suggests that environmental perturbations unique to the PWS region, including oil spills and disease outbreaks, affected the fitness of resident fish, and shifted their population genetic trajectory.

## Introduction

Rapid and radical environmental changes caused by human activities define the Anthropocene, and these pose major challenges for the resilience of wild species. When physiological or behavioral defenses fail, populations tend to decline rapidly, often to much diminished levels (or extinction) unless rescued by management interventions or adaptive evolution. Because human-induced environmental change is often multi-faceted, the specific drivers of population decline can be difficult to discern and therefore to predict. Similarly, when management interventions are ineffective, the reasons for lack of demographic recovery can be difficult to discern. Genetic data collected across space and time can illuminate the causes of demographic declines especially if mortality was selective (e.g., De Wit et al. 2014), and reveal impacts on population genetic diversity which may in turn influence recovery and adaptive potential (Spielman et al. 2004; Exposito-Alonso et al. 2022). We examined genetic variation across space and time to elucidate the causes and consequences of a notorious population collapse – the Prince William Sound (PWS) stock of Pacific herring (*Clupea pallasii*).

Pacific herring in PWS, Alaska, collapsed in the early 1990s (Trochta et al. 2020). The fishery declined more than four-fold, from approximately 133 thousand metric tons of spawning biomass in 1988 to 30 thousand metric tons by 1993, and has yet to recover (McGowan et al. 2021; Trochta and Branch 2021). In the Spring of 1989, the *Exxon Valdez* oil tanker ran aground and spilled 11 million gallons of crude oil into PWS. The *Exxon Valdez* oil spill (EVOS) devastated local ecosystems, captured international attention, and ignited a new era of research into the ecotoxicological impacts of crude oil (Peterson et al. 2003). Though the timing of the EVOS conspicuously preceded the PWS collapse, some consider its causal role controversial because of the ∼3-year lag between events, and because of co-occurrence with other hypothesized drivers such as climate regime change, starvation, and disease (e.g., Pearson et al. 1999; Rice and Carls 2007; Thorne and Thomas 2008). Furthermore, to date, the stock shows limited signs of recovery despite decades of careful monitoring and management. The reasons for the lack of recovery have yet to come into sharp focus, likely because of complex ecological interactions and additional anthropogenic influences such as changes in predator abundance and climate change (Pearson et al. 1999, 2012; Williams and Quinn, Ii 2000; Deriso et al. 2008; Ward et al. 2017; Trochta and Branch 2021).

The geographic extent of oiling following the EVOS coupled with evident toxicity at extremely low doses makes contaminating oil a likely contributor to herring population decline in PWS. Crude oil exposure causes a diverse suite of toxic syndromes, but at the lowest doses causes cardiovascular system developmental malformations during embryogenesis (Cherr et al. 2017). These impacts are caused by the polycyclic aromatic hydrocarbon (PAH) fraction of crude oil. Heart malformations emerge at parts-per-trillion (pptr) PAH concentrations (Incardona et al. 2023). Developmental impacts of low-level exposures may be so subtle as to not manifest until later in life (Hicken et al. 2011). Subtle early-life and late-life exposures threaten fitness (Heintz et al. 2000b; Schlenker et al. 2022; Park et al. 2025a) and can propagate across multiple generations (Sun et al. 2015; Wang et al. 2016; Bautista and Burggren 2019; Jasperse et al. 2019; Philibert et al. 2019; Park et al. 2025b). Herring were likely impacted by oil toxicity over multiple years because 1) the EVOS coincided with herring spawning (Carls et al. 2002), 2) developing fish are exquisitely sensitive to oil toxicity, 3) herring spawning and early life development happen in nearshore environments, which were heavily oiled, and 4) oil lingered in nearshore spawning habitats over subsequent years (Short et al. 2004, 2006, 2007). Natural Resource Damage Assessments estimated that up to 50% of embryos could have been negatively affected, citing a parts-per-billion threshold for toxicity (Brown et al. 1996; Carls et al. 2002). However, given updated toxicity thresholds in the pptr range (Incardona et al. 2023), the extent of impact on PWS herring fitness has likely been underestimated.

Disease outbreaks (viral hemorrhagic septicemia virus – VHSV – and the pathogen *Ichthyophonus hoferi*) swept through the PWS herring stock, but not others in the early 1990s (Marty et al. 2003), and the lack of recovery of PWS fish is at least partially attributable to recurring outbreaks (Marty et al. 2010). This is conspicuous because other nearby stocks were in demographic synchrony with PWS prior to the collapse (Rice and Carls 2007), and disease agents had been present but did not cause epizootics in other herring populations. This suggests that conditions unique to PWS in the early 1990s contributed to the disease outbreaks. It is plausible that PWS fish exposed to oil during early life in 1989 (and subsequent years) developed compromised performance such that they were susceptible to disease when recruiting into the fishery at maturity - three years later - coincident with the timing of collapse. It is also plausible that strong natural selection favoring reduced sensitivity to PAH toxicity arose at the cost of immune function. For example, urban populations of *Fundulus* killifish have rapidly evolved resistance to contaminating PAHs, where genetic variants that disabled aryl hydrocarbon receptor (AHR) signaling were favored by natural selection (Reid et al. 2016; Oziolor et al. 2019; Miller et al. 2024). Since normal AHR signaling interacts with immune system signaling, impairments in AHR function that are adaptive for toxicity resistance could be accompanied by impaired immune function. Indeed, evidence for subsequent (perhaps compensatory) selection on immune system genes was detected in PAH-adapted killifish (Reid et al. 2016). Chemical poisons and pathogens can act as powerful selective agents.

The following set of non-mutually exclusive hypotheses regarding the causes and consequences of the PWS herring collapse was formulated based on the unique history of ecological perturbations in PWS, and on evolutionary outcomes following PAH pollution exposure in other eco/evolutionary model systems (e.g., killifish): 1) Exposure to the EVOS was consequential for the fitness of PWS herring. If this is true, we predict signatures of natural selection in genes that mediate PAH toxicity, such as those that control AHR signaling and those that regulate calcium and potassium flux, in developing cardiomyocytes. 2) Disease outbreaks in the early 1990s were consequential, and remain consequential, for the fitness of PWS herring. If this is true, we predict persistent signatures of natural selection in genes that mediate disease and immunity. 3) Oil exposure acted additively or synergistically with disease outbreaks to compromise the fitness of PWS herring. If this is true, we predict signatures of natural selection in genes that mediate PAH toxicity and disease/immunity. 4) The demographic collapse eroded genetic diversity, thereby imposing limits on the pace of evolutionary rescue. If this is true, we predict that genetic diversity and effective population size declined following the collapse. To test these hypotheses, we tracked genome-wide genetic variation across time, spanning the period immediately after the spill but preceding the collapse (1991) and across three subsequent decades to 2017. We compared genetic change through time between the PWS population and two reference Alaska populations that did not experience oil spills, epizootics, or population collapses. Because natural selection tends to sort among standing genetic variation affecting phenotypes with polygenetic architecture, traditional selection scan methods are often insensitive for identifying the genomic footprints of natural selection. We, therefore, used sensitive temporal covariance methods to detect subtle signatures of genome-wide polygenic natural selection (Buffalo and Coop 2019, 2020) and tested for signatures that distinguish the recent evolutionary trajectory of the PWS population.

## Results

We sequenced whole genomes of 686 individual Pacific herring from three Alaskan populations over three decades using low-coverage whole genome sequencing with an average sequence coverage depth per individual of 1.061 (Fig. 1, Table S1). Three populations comprised two from the Gulf of Alaska, including the focal PWS population and the Sitka Sound (SS) reference population located approximately 780 km southeast of PWS, and a third population (second reference population) from the Bering Sea at Togiak Bay (TB) located 770 km west of PWS across the Alaska Peninsula. Population genetic data were collected over four time points at PWS and TB from 1991 to 2017, and over three time points at SS from 1996 to 2017. All sequence reads were mapped to the reference Atlantic herring (*Clupea harengus*) genome (Pettersson et al. 2019). Mapped data were filtered to include loci with the genotyping level of >0.5 (loci that existed in at least 50% of individuals) for each population and year combination. After further filtering, a total of 351,821 variable loci were included in the subsequent analyses.

**Figure 1:**
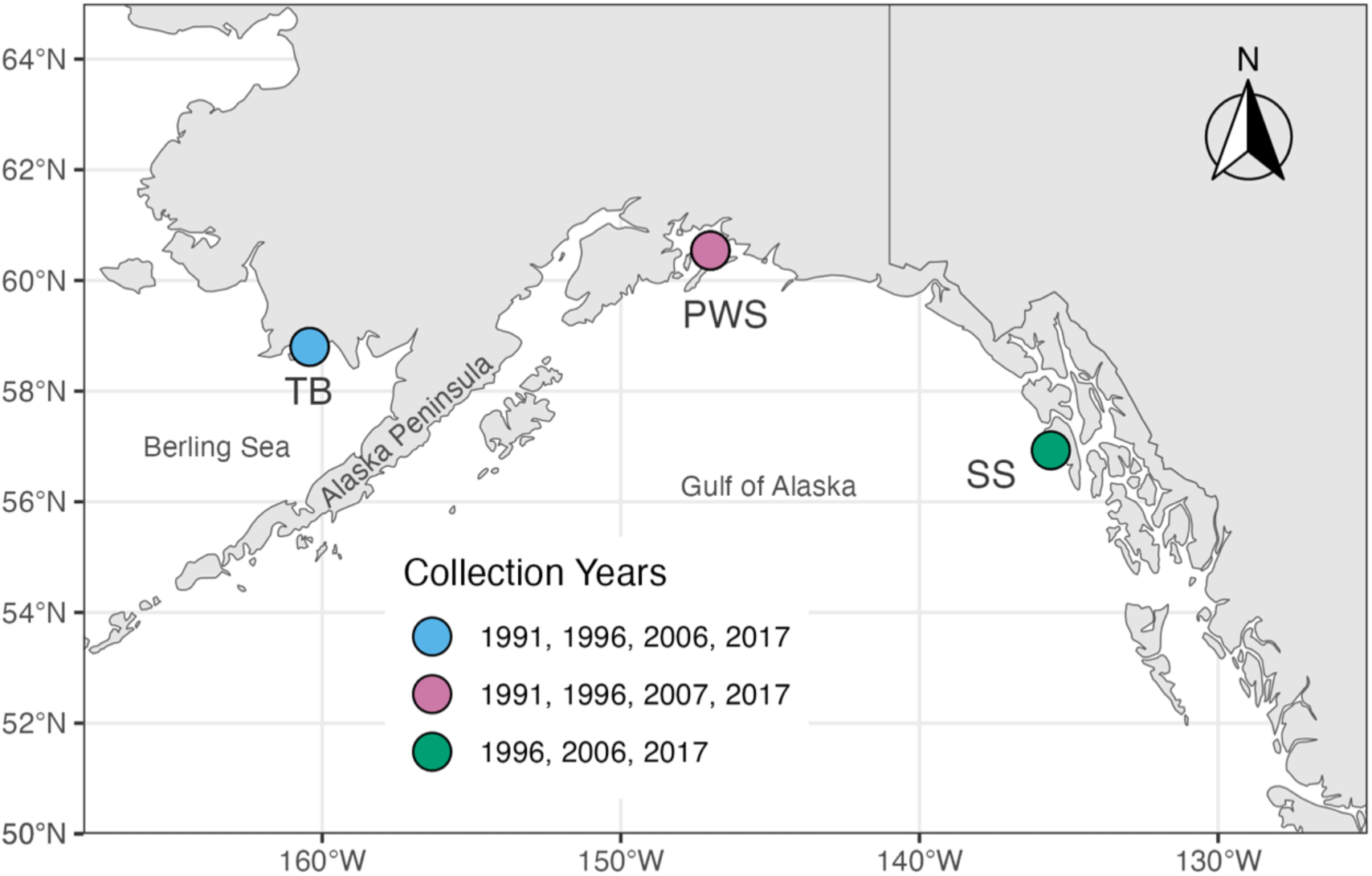
Collection sites and years of Pacific herring samples used in this study. Fish from Prince William Sound (PWS) and Togiak Bay (TB) were collected over four time points, while those from Sitka Sound (SS) were collected from three time points.

### Stable population structure in Alaska over time and space

To characterize spatial and temporal population structure, we conducted principal component analysis (PCA), model-based admixture analysis, and estimation of F_ST_ values using the ANGSD pipeline developed for low-coverage whole genome sequencing data (Korneliussen et al., 2014). PCA and admixture analysis showed clear population genetic differentiation between the Bering Sea population (TB) and the two Gulf of Alaska populations (PWS and SS) with much shared genetic variation within the Gulf of Alaska (Fig. 2) as previously reported (O’Connell et al. 1998; Wildes et al. 2018). This pattern was consistent over time and across chromosomes (Fig. S1), indicating that the Alaska Peninsula is a substantial geographical barrier to Pacific herring dispersal. Pairwise F_ST_ values are aligned with PCA and admixture results: F_ST_ between TB and PWS/SS were consistently an order of magnitude higher (0.125 - 0.170) than those between PWS and SS (0.0051 - 0.0120) (Fig. 3a). Temporal F_ST_ values within each population were stable in all three populations, ranging from 0.0066 to 0.010 in PWS, 0.0090 to 0.0092 in SS, and 0.0069 to 0.0079 in TB (Fig. 3b). Absolute genetic sequence divergence (Dxy) showed a similar stable pattern over time within populations (Fig. S2).

**Figure 2:**
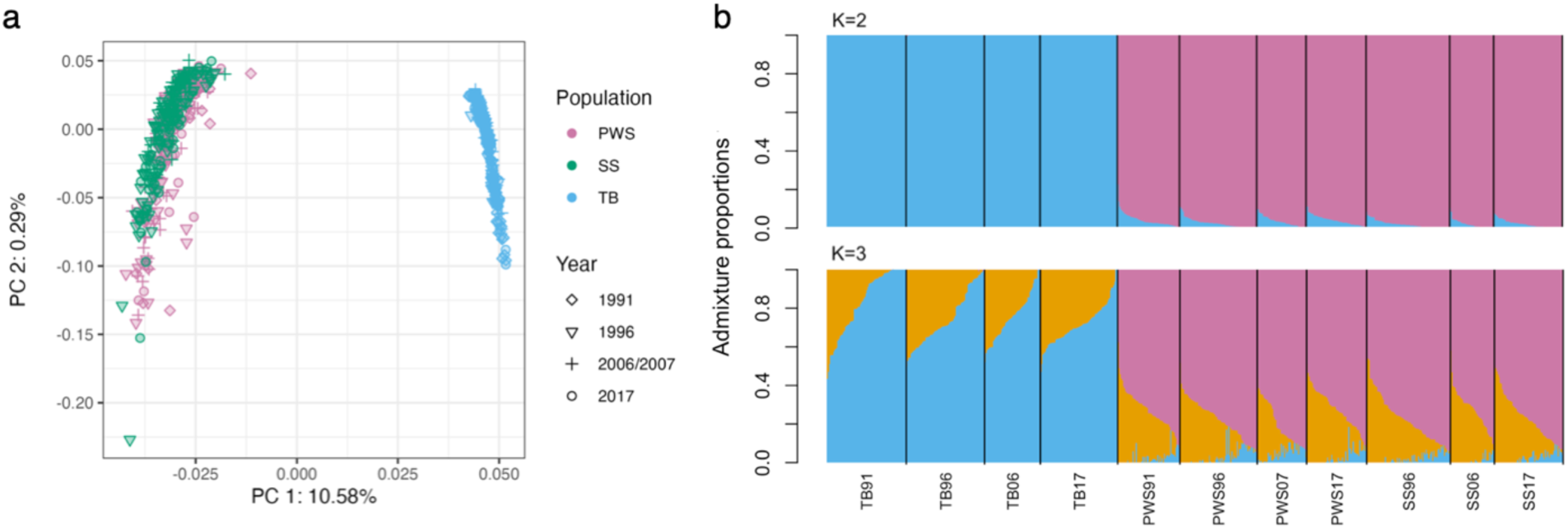
Population structure of Alaskan Pacific herring over space and time. a) PCA, revealing a clear separation between PWS/SS and TB, and b) results of Admixture for K=2 and K=3.

**Figure 3:**
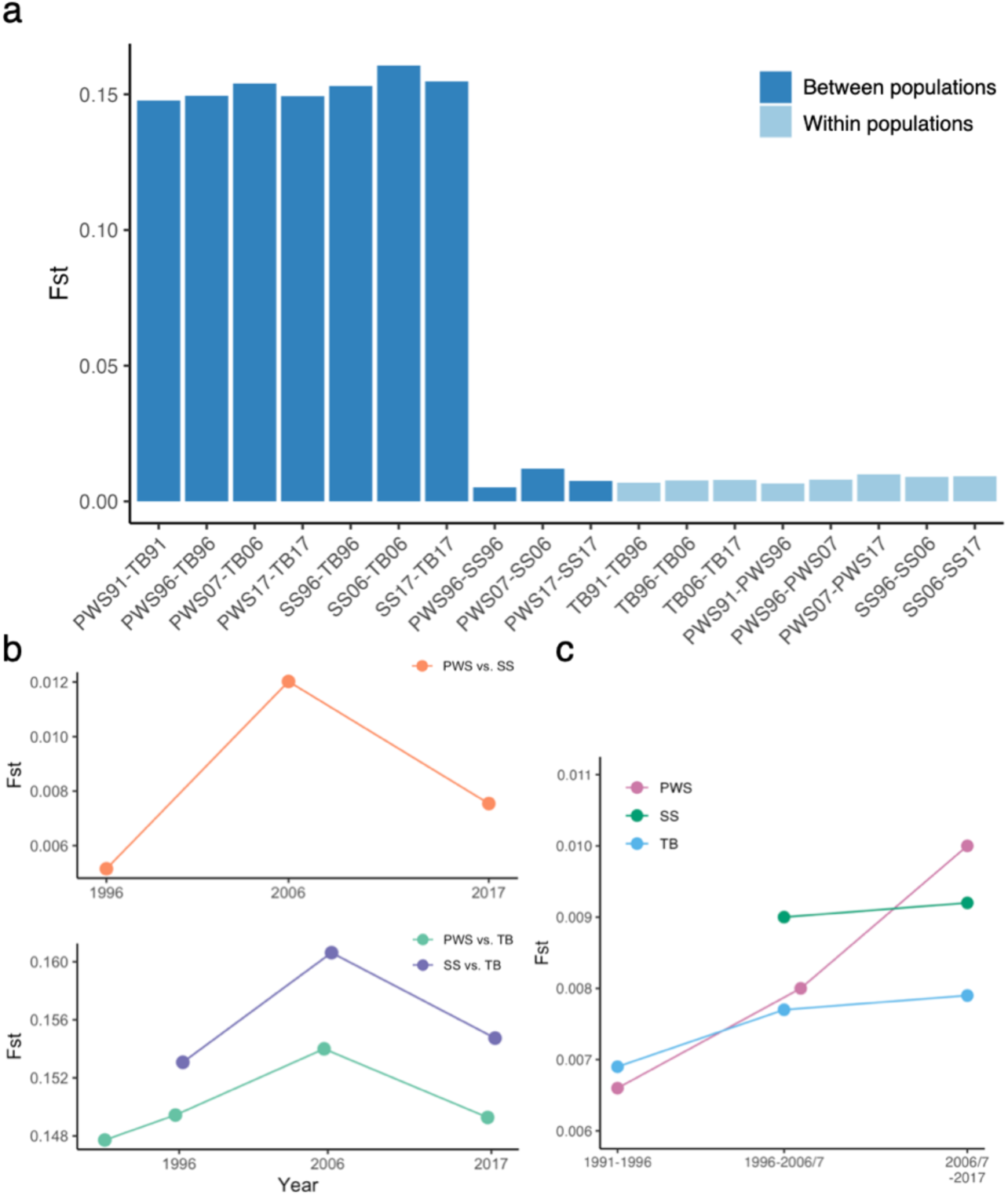
Temporal changes of F_ST_. (a) Pairwise F_ST_ between populations (in the same year) and within populations (over time), (b) F_ST_ change over time between a pair of populations, and (c) F_ST_ change over time within each population.

To evaluate whether the observed small genome-wide average F_ST_ values (0.0051-0.0120) represented significant genetic structuring between PWS and SS populations, as well as over time within each population, we performed 100 bootstraps of samples randomly assigned to a population. All measured F_ST_ values were significantly greater than random (*P*-values 0.0096 - 0.019, Fig. S3), indicating that despite much shared genetic variation, especially between fish from PWS and SS, they comprise genetically distinguishable groups. The results also revealed that small F_ST_ values between time points were significantly greater than random, highlighting the influence of genetic drift consistent with the stochastic nature of recruitment in Pacific herring (Trochta et al. 2020).

We also conducted a population-based spatiotemporal PCA using the centered and standardized population allele frequencies, showing how each population changed over time (Fig. S4a). The results confirmed that population structure was overall stable through time, with the TB population clearly differentiated from the Gulf of Alaska populations (PWS and SS). Excluding the TB population (Fig. S4b) from the PCA shows parallel allele frequency shifts in both PWS and SS in 2006/2007, for which we do not have explanations at this time.

### Little change in genetic diversity over time associated with population collapse

To determine if genetic erosion was a consequence of the population collapse in PWS, we assessed the levels of population genetic diversity (mean nucleotide diversity [π] and heterozygosity [*He*]) across space and time. Since both π and *He* tend to be correlated with read depth in low-coverage sequencing data (Li et al. 2011; Martin et al. 2021), we calculated these metrics using the single-read sampling method using ANGSD, which removed the significant correlation. We observed little variation in π and *He* across space and time, and no variation that distinguished PWS fish from other populations (Fig. 4). Despite the collapse of the PWS population and its lack of recovery, we found no evidence of genetic erosion across the subsequent three decades based on the metrics we assessed. However, these metrics can be insensitive to sudden population decline (Allendorf 1986).

**Figure 4:**
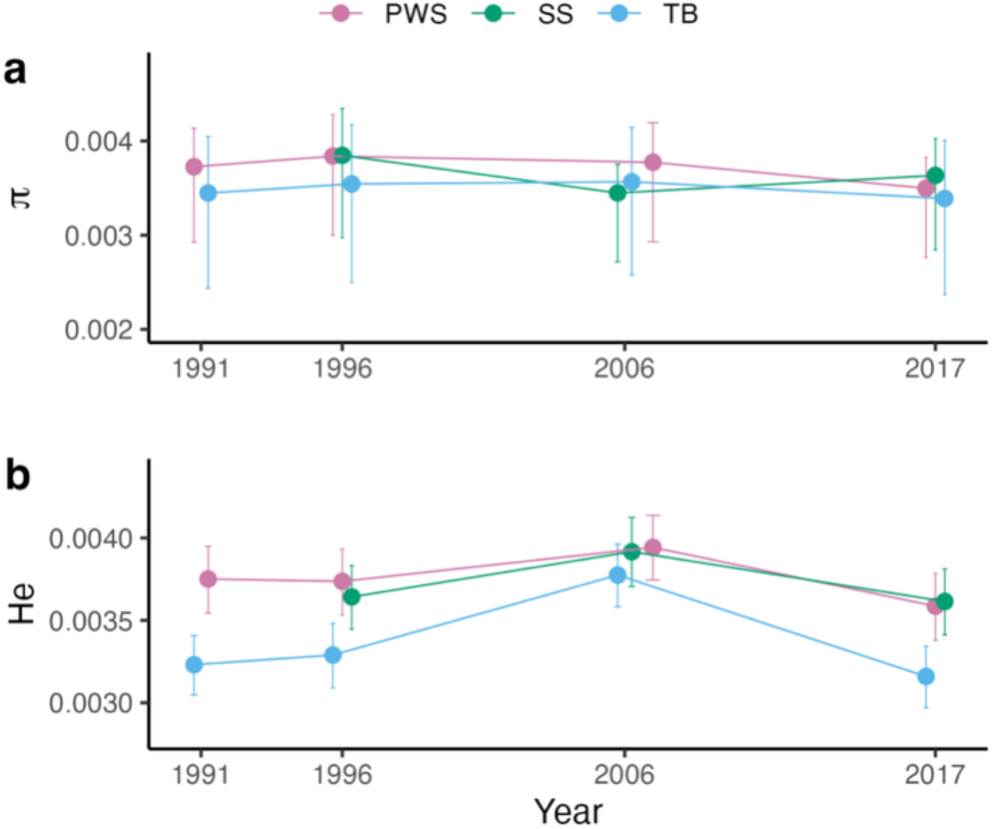
Genetic diversity variation over time in Alaskan herring populations. a) Genome-wide average nucleotide diversity. b) genome-wide average heterozygosity. Whiskers represent lower and upper quantiles.

### Estimated effective population size and recent demographic history of three populations

We estimated the contemporary effective population size (Ne) from allele frequency changes (Jorde and Ryman 2007) using NeEstimator 2.1 (Do et al. 2014) based on the longest time intervals (1991-2017 for PWS and TB, 1996-2017 for SS). Estimates of contemporary effective population sizes were small: 234 for PWS, 198 for TB, and 229 for SS population. We also estimated Ne for each time period, which generated even smaller Ne estimates (40 – 85, Fig.5A). The Ne values estimated using this method may not be accurate due to the small sample size relative to the census population size and the relatively short generation intervals (Marandel et al. 2019), especially for separate time periods.

**Figure 5:**
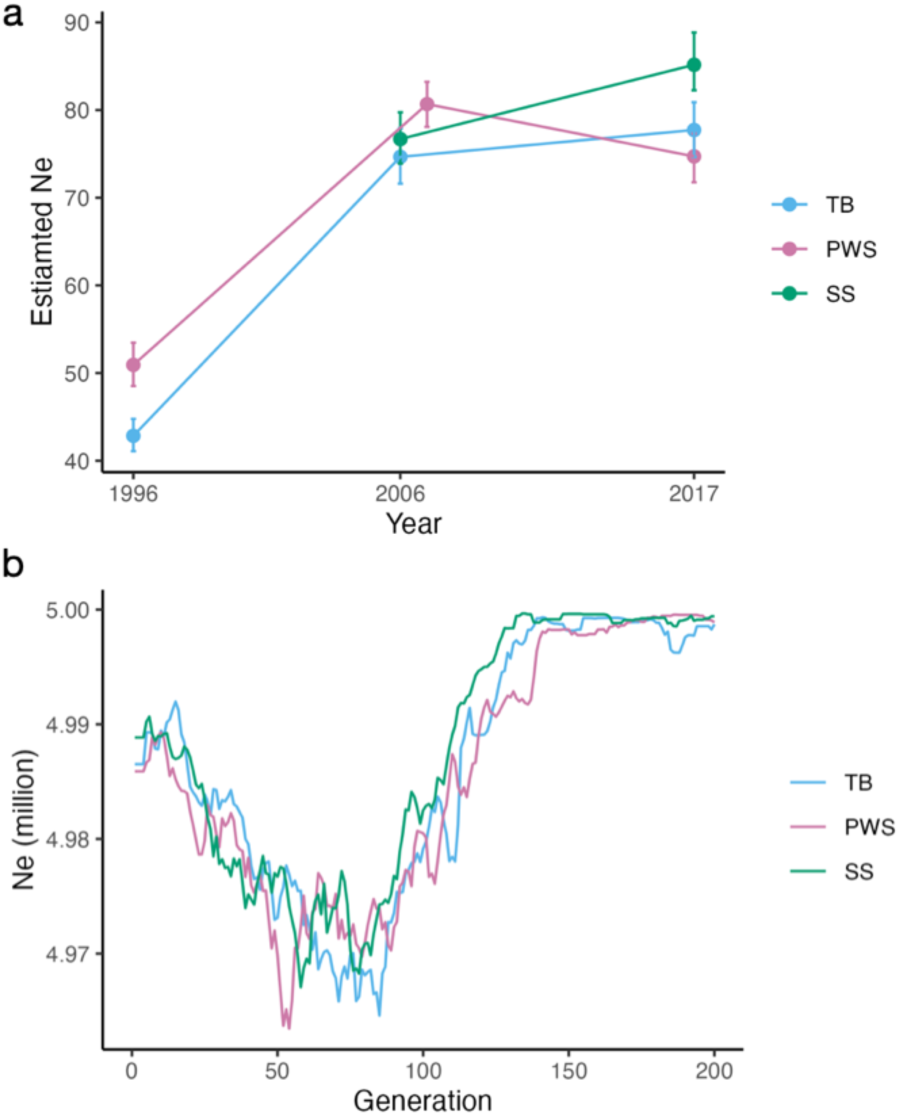
Estimated effective population sizes (*Ne*) over time in three Alaska herring populations. a) *Ne* estimated from short-term allele frequency changes based on the Jorde and Ryman method (Jorde and Ryman 2007). Whiskers represent 95% confidence intervals. b) Population demographic history for 200 generations estimated using Gone (Santiago et al. 2020).

However, the patterns of variation in Ne that we observed within and between populations were consistent across the different Ne estimation methods that we tested (NeEstimator [Do et al. 2014], poolSeq [Taus et al. 2017], *cvtk* program, [Buffalo and Coop 2020]): Both PWS and TB populations showed an increase in Ne in Period 1 (1991-2006/7), and all populations showed little change in values afterward (Period 2), with PWS being the only one with a slight declining trend in Period 2 (Fig. 5A). The observed small Ne values across different populations also support the stochastic nature of herring recruitment with few individuals likely contributing to the next generation. No decline in PWS Ne is consistent with our findings of no erosion of genetic diversity, which would be unlikely to be detected given the very small Ne.

The demographic history of herring populations over a longer time period (up to 200 generations) was estimated with the 2017 data using GONE (Santiago et al. 2020) (Fig. 5B). All three populations showed similar trends, including a decline in Ne from about 125 generations ago (about 750 years ago) followed by recovery starting approximately 50 generations ago (about 300 years ago). The results from GONE showing correlated long-term demographic changes across all three populations are consistent with observed demographic coupling of contemporary herring populations in Alaska (Trochta et al. 2020).

### Traditional selection scans do not detect signatures of large selective sweeps

Because the PWS herring population experienced multiple environmental perturbations (oil exposure and epidemic), we conducted genome-wide scans to screen for loci that may have been responsive to natural selection. No significant signatures of natural selection were identified from PC-based genome-wide selection scans between years within populations, nor between population pairs within the same period (Fig. S5). Similarly, F_ST_ based genome scans using a null model (through shuffling loci to create null distributions, Pinsky et al. 2021) also did not detect significant (adjusted *P*-value <0.05) signatures of natural selection (Fig. S6).

### Temporal covariances show significant signatures of genome-wide polygenic adaptation in PWS only, associated with heart development and immune genes

Since scans for F_ST_ outliers did not detect the influence of natural selection, we tested for more subtle signatures of genome-wide polygenic adaptation by examining patterns of temporal covariance in allele frequencies using *cvtk* (https://github.com/vsbuffalo/cvtkpy). This approach can detect polygenic selection over short timescales (Buffalo and Coop, 2019, 2020), because indirect selection at linked sites creates covariances in allele frequency changes at different time points that cannot be generated by random drift alone. PWS and TB populations were sampled at four timepoints which enabled three temporal covariance measurements (Period 1: 1991-1996 vs. 1996-2006, Period 2: 1996-2006 vs. 2006-2017, Period 3: 1991-1996 vs. 2006-2017) while the SS population was sampled at three timepoints such that temporal covariance was estimated for the period 1996-2006 vs. 2006-2017 only (Period 2). The results revealed signatures of genome-wide positive selection (polygenic adaptation) in the PWS population, while these signals were not observed in TB nor SS populations (Fig. 6). Indeed, the PWS population in Period 1 was the only one with a positive temporal covariance estimate (0.00072 ± 0.00022). This is consistent with the PWS herring population subjected to selective pressures that were unique to that region over that time frame. All temporal covariance estimates for reference populations were negative (ranging from −0.00324 to −0.000326), indicating reversals in selection across time.

**Figure 6:**
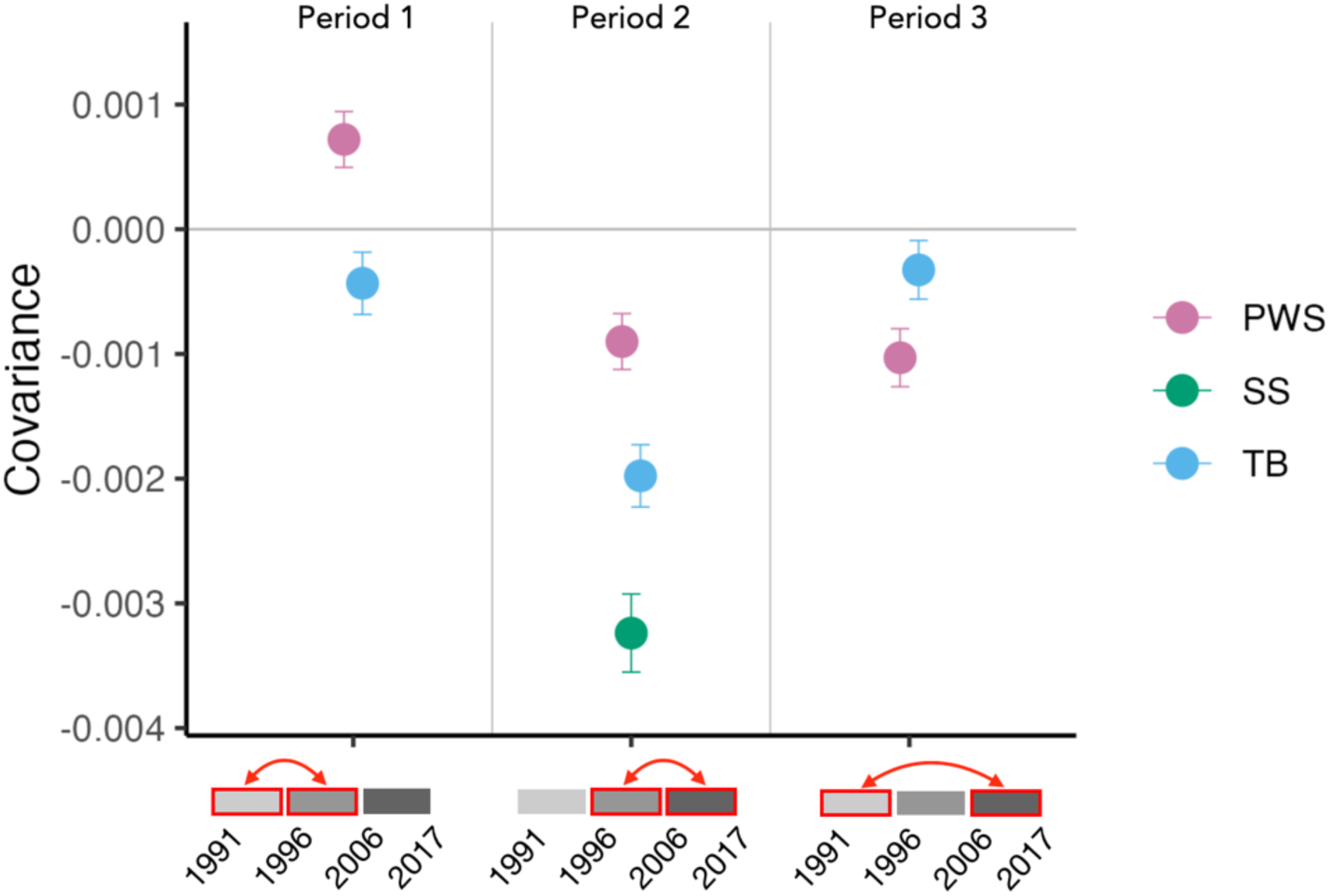
Covariances of allele frequency changes in three herring populations. The mean genome-wide covariance calculated from *cvtk* for each time period (red boxes) for each population. Whiskers represent 95% confidence intervals.

Since the *cvtk* program was established to detect linked selection using allele frequency covariances from populations with non-overlapping generations, we assessed how overlapping generations might have impacted the observed patterns of temporal covariances using simulations in SLiM (Haller and Messer, 2019). The generation time of Pacific herring in the Pacific north east is estimated to be 4-6 years, with the age of first reproduction around 2-3 years (Cleary et al., 2010). Under the neutral selection scenario, both overlapping and non-overlapping generation models produced highly similar covariance patterns (Fig. S7), such that we conclude that *cvtk* can be appropriately applied for species with overlapping generations as long as they are not sampled every year.

We identified candidate genes subject to natural selection in the PWS population from Period 1 using the genomic windows (100kb) with the highest (top 1%) temporal covariance measures (Fig. 7A). Among the 289 protein-coding genes identified within these regions (Appendix S1A), the two most highly enriched GO terms were *C-C chemokine receptor activity* (related to immune response) and *cardiac muscle tissue development* (Fig. S8, see Appendix S1B for all enriched GO terms). Other genes involved in immune function and heart development were also among the candidate genes (Table 1). Multiple alleles within the large (25 kb) immune gene *C-C chemokine receptor type 6* (ccr6a), and the pollution-responsive heart development gene *aryl hydrocarbon receptor nuclear translocator* (arnt), show parallel patterns of positive temporal covariance in Period 1 in the PWS population (Fig. 7b, 7c).

**Figure 7:**
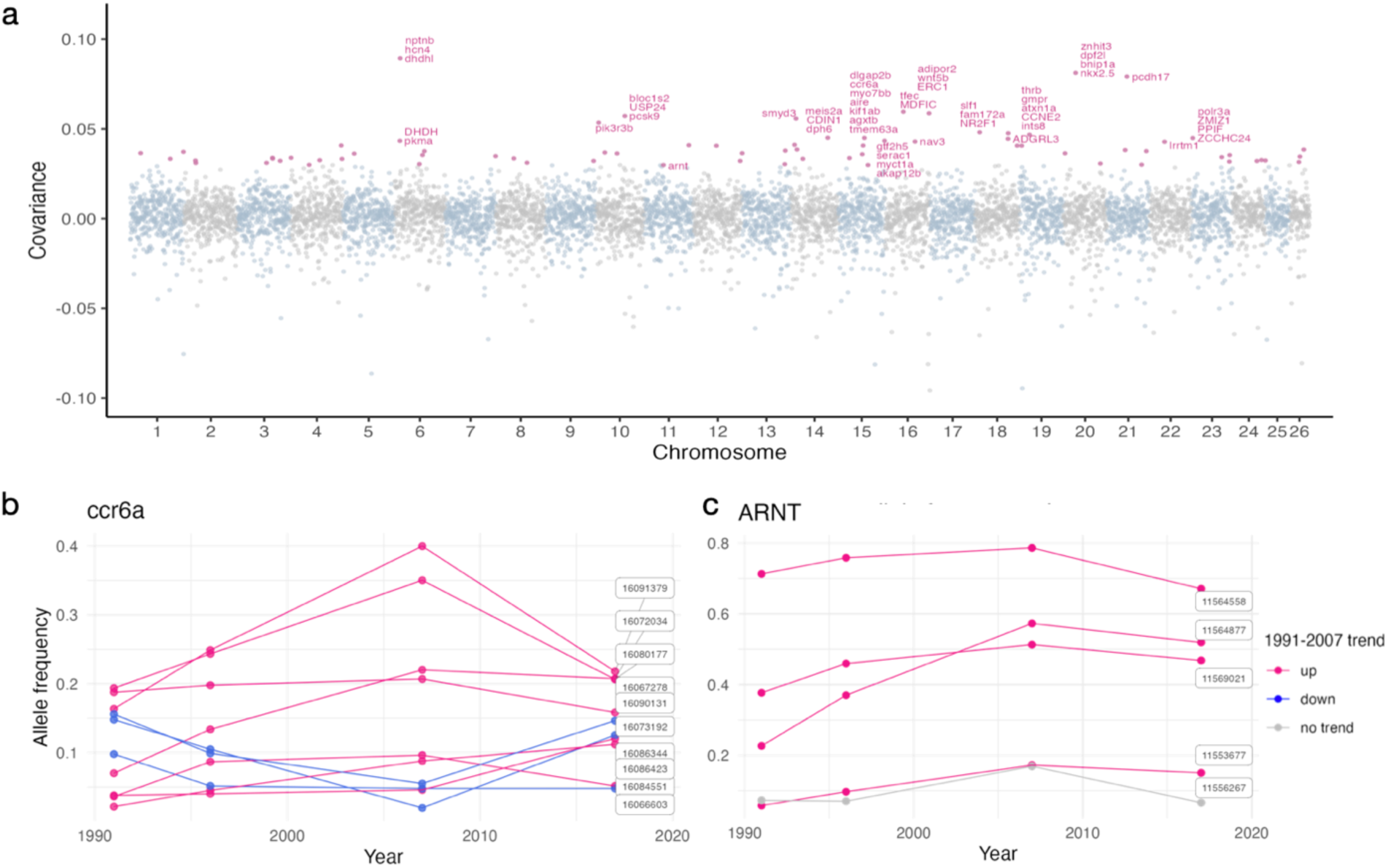
Genes identified in the high covariance outlier regions identified from the temporal covariance (*cvtk*) analysis in the PWS population during Period 1 (1991-2007). a) Points represent mean covariance for 100kbp windows. Points in magenta were outliers (top 1%), and genes identified in the outlier windows with covariance >0.4 are labeled. Allele frequency changes at individual loci that belonged to ccr6a (b) and ARNT (c) genes. For ccr6a, only the loci that showed an upward (magenta) or downward (blue) allele frequency trend in Period 1 are shown.

**Table 1:** List of candidate genes involved in immune function and heart development identified from the outlier genomic regions (potentially selected) from the covariance analysis in the PWS population from Period 1 (1991-1996 vs. 1996-2007).

| Gene symbol | Gene name | Function |
| --- | --- | --- |
| **hcn4** | hyperpolarization activated cyclic nucleotide-gated potassium channel 4 | Heart contraction |
| **cmlc1** | cardiac myosin light chain-1 | Heart development |
| **nkx2.5** | NK2 homeobox 5 | Heart development |
| **opa1** | OPA1 mitochondrial dynamin like GTPase | Heart development |
| **sgcd** | sarcoglycan delta | Heart development |
| **smarca4a** | SWI/SNF related BAF chromatin remodeling complex subunit ATPase 4a | Heart development |
| **synpo2la** | synaptopodin 2-like a | Heart development |
| **aire** | autoimmune regulator | Immune related |
| **bloc1s2** | biogenesis of lysosomal organelles complex-1, subunit 2 | Immune related |
| **bnip1a** | BCL2 interacting protein 1 | Immune related |
| **C3** | complement C3 | Immune related |
| **ccr6a** | C-C chemokine receptor type 6-like | Immune related |
| **CDIN1** | chromosome unknown open reading frame, human C15orf41 | Immune related |
| **gfi1ab** | growth factor independent 1A transcription repressor b | Immune related |
| **MDFIC** | MyoD family inhibitor domain containing | Immune related |
| **mettl22** | methyltransferase like 22 | Immune related |
| **myct1a** | myc target 1a | Immune related |
| **polr3a** | polymerase (RNA) III (DNA directed) polypeptide A | Immune related |
| **slc25a38a** | solute carrier family 25 member 38a | Immune related |
| **wnt5b** | wingless-type MMTV integration site family, member 5b | Immune related |

## Discussion

We detected genetic structuring across space and time in Alaska populations of Pacific herring. Genetic differentiation was small but statistically significant and temporally stable between the two neighboring Gulf of Alaska PWS and SS populations (F_ST_ ∼ 0.01). PWS and SS stocks showed demographic coupling over time, at least prior to the 1990s (Rice and Carls 2007; Pearson et al. 2012), likely because they experienced similar environmental conditions within the Gulf of Alaska rather than because they were panmictic. The structuring that we detected between PWS and SS may be noted by resource managers as they consider their management as distinct stocks (Hay et al. 2001). In contrast to the subtle genetic differences between populations within the Gulf of Alaska, we detected substantial genetic differentiation between fish from the Gulf of Alaska and the Bering Sea (F_ST_ ∼ 0.15). This is consistent with reports from other herring studies (e.g., Grant and Utter 1984; O’Connell et al. 1998; Liu et al. 2012; Timm et al. 2025) and other marine species (e.g., Talbot et al. 2016) indicating that the Alaska Peninsula provides a significant barrier to marine dispersal. Multiple estimates indicate very small contemporary effective population size (Ne) for all Alaskan herring populations at all timepoints, especially compared to the very large census population size (Nc) for Pacific herring populations (>hundreds of millions fish, Rice and Carls 2007). Although large sample sizes may be necessary to accurately estimate Ne in species with very small Ne (Waples 2016), we conclude that the ratio of Ne to Nc is extremely small in Pacific herring. This small Ne is consistent with sweepstakes-style reproduction strategies (high variance in reproductive success between individuals), which appear to be common for many highly fecund marine species (Hedgecock and Pudovkin 2011), resulting in Ne to Nc ratios ∼10^-3^ to 10^-6^ or even smaller (Nunney 1993; Waples 2016; Buffalo 2021).

Our genetic diversity estimates appeared to be stable over time in all Alaska herring populations, which was inconsistent with the hypothesis that the PWS population collapse caused an erosion of genetic diversity. This may be because very large population bottlenecks are necessary to cause detectable declines in nucleotide diversity, and the ∼80% decline observed in PWS herring - occurring between our first and second sampling timepoints - may have been insufficient to cause such a change (Osmond and Coop 2020). Alternatively, too few generations may have passed since the population collapse for a reduction in diversity to manifest, as detectable losses can lag by hundreds to thousands of generations (Allendorf 1986; Petit-Marty et al. 2022). Gene flow among Gulf of Alaska populations may also buffer against diversity loss (Rice and Carls 2007). However, our temporal data enabled more sensitive short-term estimates of contemporary effective population size (Ne), which consistently showed no significant decline. Collectively, these findings suggest that, up to our latest sampling in 2017, the erosion of genetic diversity was not a major factor impeding recovery of the PWS herring population.

Temporal patterns of genetic change in the PWS population revealed signals of positive selection and polygenic adaptation during Period 1 (1991–2007), coinciding with major ecological disturbances such as the EVOS and viral epidemics that distinguished PWS from other regions. The identity of candidate genes implicated in these adaptive responses offers clues to the ecological drivers of fitness challenges experienced by PWS fish. We observed significant enrichment of genes involved in immune function and heart development. Particularly, this enrichment of genes involved in immune function (e.g., *C-C chemokine receptor activity*) is conspicuous since this was the period over which VHSV swept through the PWS population. Chemokines and C-C chemokine receptors play major roles in immune system signaling, inflammatory reactions, and innate immunity, and are known to be specifically induced by exposure to VHSV in fish (Sanchez et al. 2007; Chaves-Pozo et al. 2010; Montero et al. 2011; Aquilino et al. 2014; Castro et al. 2014; Kim et al. 2018). Immune system genes showing signals of polygenic adaptation in the PWS population (Period 1) include C-C chemokine receptor type 6-(ccr6a), which has a key role in adaptive immunity (Dieu et al. 1998), *complement component 3* (C3), which contributes to regulation of innate immunity (De Bruijn and Fey 1985), *autoimmune regulator* (aire) which enables the immune system to distinguish host from foreign cells (Anderson and Su 2011), *BCL2 interacting protein1* (bnip1a) which protects from virally induced cell death (Liu et al. 2024), and *MyoD family inhibitor domain containing protein* (MDFIC) which regulates viral genome expression (Byrne et al. 2022).

Enrichment of genes involved in cardiac muscle development in PWS in Period 1 is also conspicuous because it is coincident with PWS herring exposure to EVOS crude oil during development, which canonically perturbs the development of heart structure and function with significant impacts on performance and fitness (Heintz et al. 2000a; Hicken et al. 2011). Cardiovascular developmental toxicity caused by PAH exposure is mediated partially through perturbed aryl hydrocarbon receptor (AHR) signaling and partially through perturbation of calcium and potassium fluxes that regulate excitation-contraction coupling in developing cardiomyocytes (Brette et al. 2014). Evolved resistance to PAH toxicity in killifish is associated with altered AHR signaling and with signatures of selection in AHR pathway genes and in pathways that regulate intracellular calcium and potassium (Whitehead et al. 2012; Reid et al. 2016; Miller et al. 2024). Outlier genes in PWS Period 1 include arnt, which regulates PAH-induced AHR signaling that causes cardiovascular system defects. Variants of arnt appear subject to strong natural selection in PAH-adapted killifish populations (Reid et al. 2016; Oziolor et al. 2019).

Accompanying arnt, other outlier genes in PWS Period 1 involved in fish cardiac muscle tissue development and function include *potassium/sodium hyperpolarization-activated cyclic nucleotide-gated channel 4* (hcn4), which regulates potassium flux in the heart cells that regulate contraction and rhythm (Jou et al. 2017), *PR domain zinc finger protein 16 isoform X7* (prdm16), which affects excitation-contraction coupling in developing cardiomyocytes (Arndt et al. 2013), *synaptopodin 2-like protein isoform X1* (synpo2la) and *cardiac myosin light chain-1* (cmlc1), which are both expressed in developing cardiomyocytes and are important for sarcomere structure and heart contractility (Meder et al. 2009; Beqqali et al. 2010), *homeobox protein Nkx-2.5* (nkx2.5) and *brahma protein-like protein 1* (smarca4a) which interact to regulate cardiac differentiation and heart tube elongation (Targoff et al. 2008; Takeuchi et al. 2011), *nuclear receptor subfamily 2 group F member 1* (nr2f1) which regulates atrium development (Duong et al. 2018), *delta-sarcoglycan isoform X1* (sgcd) which affects cardiac muscle differentiation and symmetry (Cheng et al. 2006), and *protein naked cuticle homolog 1* (nkd1) which regulates heart symmetry (Schneider et al. 2010).

Considering that the two most highly enriched categories of genes showing signatures of polygenic adaptation in PWS Period 1 include heart development and immune system genes is consistent with our hypotheses; that the EVOS oil spill and disease epidemics – the two environmental perturbations that distinguish PWS from other populations – were consequential for PWS herring fitness during the period following the oil spill and population collapse. This is because the major mechanism of crude oil toxicity is through impairment of heart development, and cardiac development genes are subject to natural selection from PAH exposures in other model systems (e.g., Reid et al. 2016; Miller et al. 2024), and because the immune system mediates sensitivity to pathogens, and immune system genes are subject to natural selection following pathogen exposures in other model systems (e.g., Fraslin et al. 2020).

Despite signatures of selection for genes involved in immune function and pollution-sensitive heart development in PWS fish, there is little evidence for evolved divergence in sensitivity to crude oil toxicity or sensitivity to virus exposure in contemporary PWS fish compared to other populations (Hershberger et al. 2010). We consider some potential explanations for this. First, it is plausible that natural selection resulting from oil and pathogen exposures that distinguished PWS from other populations is manifest in traits that have not yet been measured. Second, perhaps selection drove allele frequency change for alleles affecting sensitivity to oil toxicity during the period after the spill, but in subsequent years, when toxicity in PWS habitats was diminishing, the selection pressure eased and allele frequency change stopped or reversed because of adaptive tradeoffs. Indeed, the pattern of positive covariance detected in early periods (Period 1) was followed by negative covariance in the final period (Period 2) in PWS (Fig. 6). Fluctuations in the directionality of natural selection, accompanied by parallel fluctuation in related traits, are likely common in natural systems.

Notably, allele frequency covariances for most other herring populations and periods were significantly negative, indicating changing directions of selection across time. Natural environments are characteristically variable, where conditions fluctuate periodically and stochastically over time. This natural variability likely imposes fluctuating selection pressures over time for most wild species. Since adaptation is often polygenic, we suspect that temporally fluctuating selection would commonly cause patterns of negative allele frequency covariances over time in many species (e.g., Machado et al. 2021). We hope future studies will further explore temporal covariance estimates using genome-wide time series data from wild populations. With more extensive datasets, we may better determine whether the evolutionary equilibrium for wild populations is due to neutral processes or is subject to ongoing, multidirectional selective pressures.

It is perhaps surprising that there has been no population recovery in recent decades, especially if adaptation was enabled by natural selection in response to environmental change, and genetic diversity was not subsequently eroded. It is plausible that new environmental stressors, such as changes in predation and climate change, have compounded the impacts of prior stressors and thereby limited the pace of recovery. It is also plausible that adaptations to the stressors of the early 1990s have enabled persistence, but are accompanied by adaptive tradeoffs that limit the pace of recovery. Another consideration is that demographic recovery lags behind adaptive changes in allele frequency (Osmond and Coop 2020), such that perhaps there has been insufficient time since adaptive allele frequency change for population recovery to have been observed. Pacific herring is one of the most intensively studied and monitored of marine fish species, such that current and ongoing research into the drivers of population dynamics are likely to further illuminate the drivers of demographic change in rapidly changing Arctic ecosystems.

## Methods

### Sample collection

Pacific herring samples were collected from three geographic locations across two ocean basins over time: PWS and TB fish were collected in four time points of 1991, 1996, 2006 (TB) or 2007(PWS), and 2017, and SS fish were collected in three time points of 1996, 2006, and 2017 (Fig. 1, Table S1).

### Library preparation, sequencing, and alignment

Whole genome sequencing of 686 individuals resulted in 6,843,679,501 raw reads that were trimmed using Cutadapt v. 1.8.3 (Martin 2011). Trimmed reads were mapped to the Atlantic herring reference genome (*C. harengus* chromosome level genome assembly Ch_v2.0.2 [Pettersson et al. 2019]; chromosomes = 26; total base pairs = 725,670,187; protein coding genes = 24,095; gene transcripts = 67,663) using the Burrows-Wheeler Alignment Tool (v. 0.7.17, Li and Durbin 2009) with an average mapping rate of 97.3% resulting in 6,663,624,351 mapped reads. We marked duplicates with Picard Tools MarkDuplicates (v. 2.7.1, broadinstitute.github.io/picard/), sorted alignments using Samtools (v.1.9, Li et al. 2009), and assessed the quality of alignments with multiQC (Ewels et al. 2016) and Picard Tools CollectWgsMetrics. Mean sequencing depth of coverage was 1.06× across all samples.

### SNP calling and genotype likelihood estimation

After mapping the reads to the reference genome, we used the Genotype Analysis Toolkit (GATK v. 4.1.4.1, Poplin et al. 2017) to call SNPs and produce phred-scaled genotype likelihoods. We generated GVCFs from sorted alignments using Haplotype Caller and followed the GATK4 Best Practice Workflow for variant calling, which uses GenomicsDBImport to merge GVCFs from multiple samples. We used GenomicsDBImport to create a database for each of the 26 chromosomes and GenotypeGVCFs to call raw variants across all samples for each chromosome. Raw variants were filtered to include biallelic SNPs at sites with Phred-scaled quality > 20, mapping quality > 30, and depth > 417 and < 2000. These filtered variants were concatenated into one VCF file containing all chromosomes with bcftools (v. 1.10.2, Li 2011) ‘concat’ function. We subset this VCF into 11 VCF files – one for each population at each timepoint. We filtered each VCF file to include sites with genotyping rate > 50%, and intersected all using the bcftools ‘isec’ function to keep only sites common across all files. Finally, we filtered out sites with a minor allele frequency < 0.05. Our final VCF included 351,820 biallelic SNPs.

### Population structure and genome-wide summary statistics

We converted genotype likelihoods from the final VCF files into BEAGLE format, then pruned the SNPs for linkage disequilibrium (--indep-pairwise 75’kb’ 5 0) using PLINK v1.90b6.24. PCAngsd (Meisner and Albrechtsen 2018) and NGSadmix (Skotte et al. 2013) were run on the pruned files to analyze population structure. We evaluated the optimal cluster number (K) using evalAdmix (Garcia-Erill and Albrechtsen 2020) and CLUMPAK (Kopelman et al. 2015). We used Analysis of Next Generation Sequencing Data software (ANGSD v. 0.931, Korneliussen et al. 2014) to calculate relative genetic divergence (F_ST_). We subset the final VCF file into 11 populations (grouped by location and year collected) and used ANGSD to calculate a folded SFS and per site allele frequencies from genotype likelihoods. We created two-dimensional SFS for each population pair and calculated F_ST_ in 50kb sliding windows with a 10kb step size. We calculated absolute genetic divergence (D_XY_) from allele frequency estimates by ANGSD using the CalcDxy.R script in ngsTools (Fumagalli et al. 2014). Since read depths and genetic diversity parameters showed a correlation, we used single read sampling (-doIBS) in ANGSD to estimate population allele frequencies and used them to measure population nucleotide diversity (π) and expected heterozygosity (He).

### Selection scans

Genome-wide selection scans were conducted using PCAngsd (Meisner and Albrechtsen 2018). F_ST_ outliers were also assessed by creating a null model for the maximum change in a region expected across the genome, following the method of Pinsky et al. (2021). Briefly, the pairs of unlinked SNP allele frequencies were shuffled across the unlinked SNP locations in the genome, and the maximum F_ST_ values over a specified window across the entire shuffled genome were recorded 1000 times. This empirical distribution of maximum F_ST_ was used to assess F_ST_ values that were significantly larger than random.

### Genome-wide linked selection using temporal covariances of allele frequency changes

We assessed the genome-wide covariance of allele frequency changes from the time-series data to determine if signals of linked selection could be detected. The temporal covariances were calculated using the *cvtk* pipeline (Buffalo and Coop 2019, 2020) for each population using 100kb windows. In addition to the genome-wide averages, we calculated the average covariances based on 100kb windows, which were used to identify the outlier regions that fell within the top 1% of the total covariances for each population within each period. Genes in the outlier regions (with 100k padding) were identified using SnpEff v. 5.1 (Cingolani et al. 2012) and the Gene Ontology enrichment analysis was carried out using ShinyGo v.0.77 (Ge et al. 2020).

### Effective population size (Ne) estimation

We estimated contemporary effective population sizes (Ne) for each population based on changes in unlinked allele frequencies over time, using the Jorde-Ryman temporal method from NeEstimator 2.1, following Pinsky et al. (2021). We also used the poolSeq package in R (available at https://github.com/ThomasTaus/poolSeq) to apply different calculation methods, including the methods of Waples (Waples 1989) and Jónás et al. (Jónás et al. 2016). Although we could not fully account for overlapping generations due to the data limitations, our sampling spanned approximately six Pacific herring generations (1991 to 2017), which likely minimized potential biases associated with overlapping generations (Waples and Yokota 2007; Waples et al. 2014). On the other hand, *Ne* numbers estimated using each sampling period (e.g., 1991-1996, 1996-2007, Fig. 5A) spanned only 1-3 herring generations, which likely strongly biased the estimates downward. We presented the results as the shapes of *Ne* changes over time did not differ from one method to another. Additionally, we estimated historical *Ne* and demographic history of three populations using GONE (Santiago et al. 2020) with the 2017 samples from each population.

## Data Availability

Raw sequencing reads for all samples used in this article are available at the NCBI SRA (PRJNA972458). Analysis scripts and associated data reported in this article are archived as a Git repository at https://github.com/awhitehe/PacificHerring_Temporal.

## Acknowledgements

This research was supported by funding from the *Exxon Valdez* Oil Spill Trustee Council. Pacific herring tissue samples were coordinated or provided by Chris Habicht, Judy Berger, Heather Hoyt, Sherri Dressel, and Katie Sechrist (Alaska Department of Fish and Game), Ben Cain (Silver Bay Seafoods, Bristol Bay/Naknek plant), and Ashley Mackenzie (United States Geological Survey). This project is part of the Herring Research and Monitoring program. These findings and conclusions presented by the author are their own and do not necessarily reflect the views or position of the *Exxon Valdez* Oil Spill Trustee Council.

## Notes

### Competing Interest Statement

The authors have declared no competing interest.

