## Supplementary material for "Evidence of positive polygenic selection following a series of environmental and demographic disasters in Pacific herring": Figures S1-S8

**Supporting Figures**

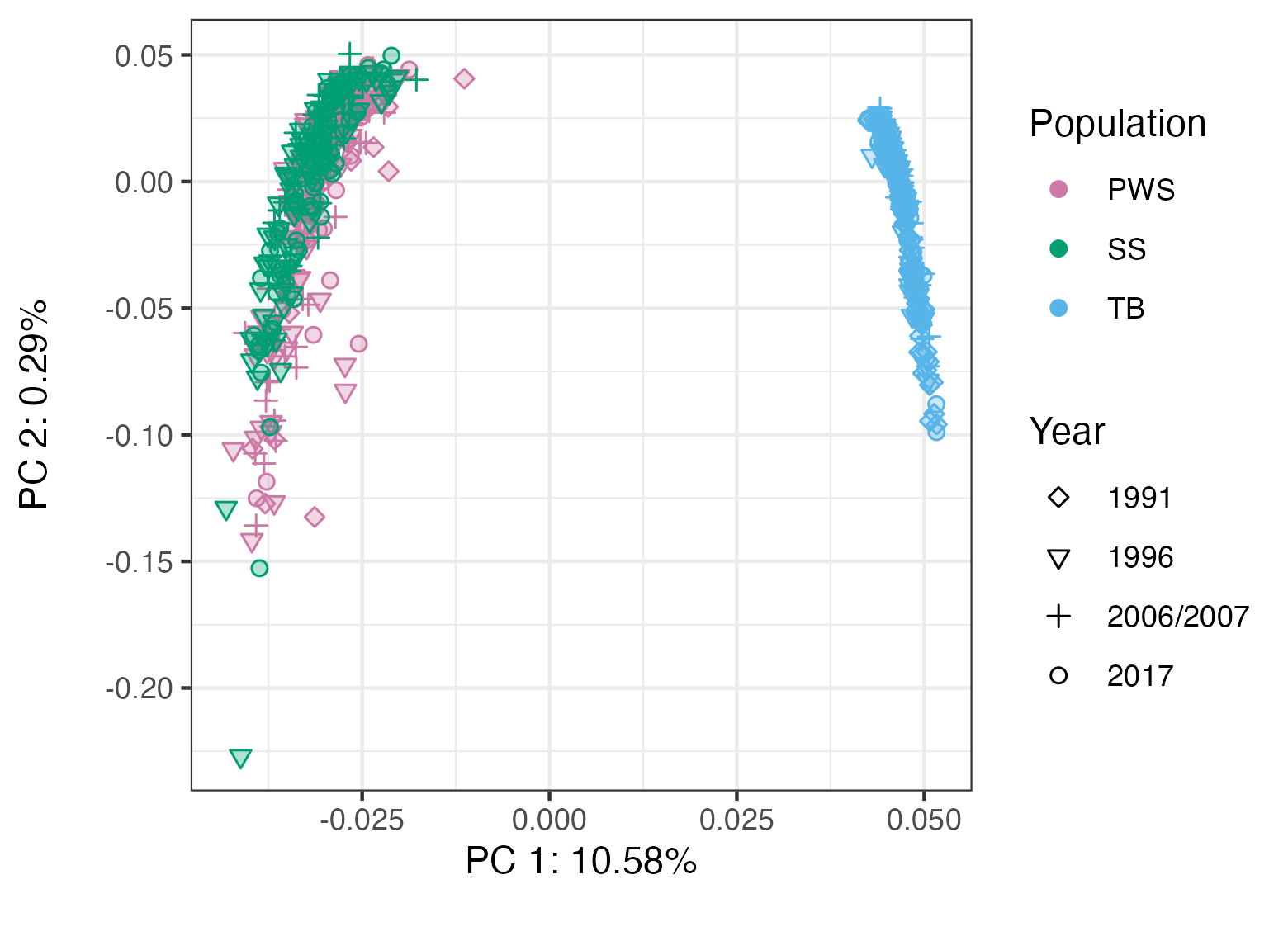

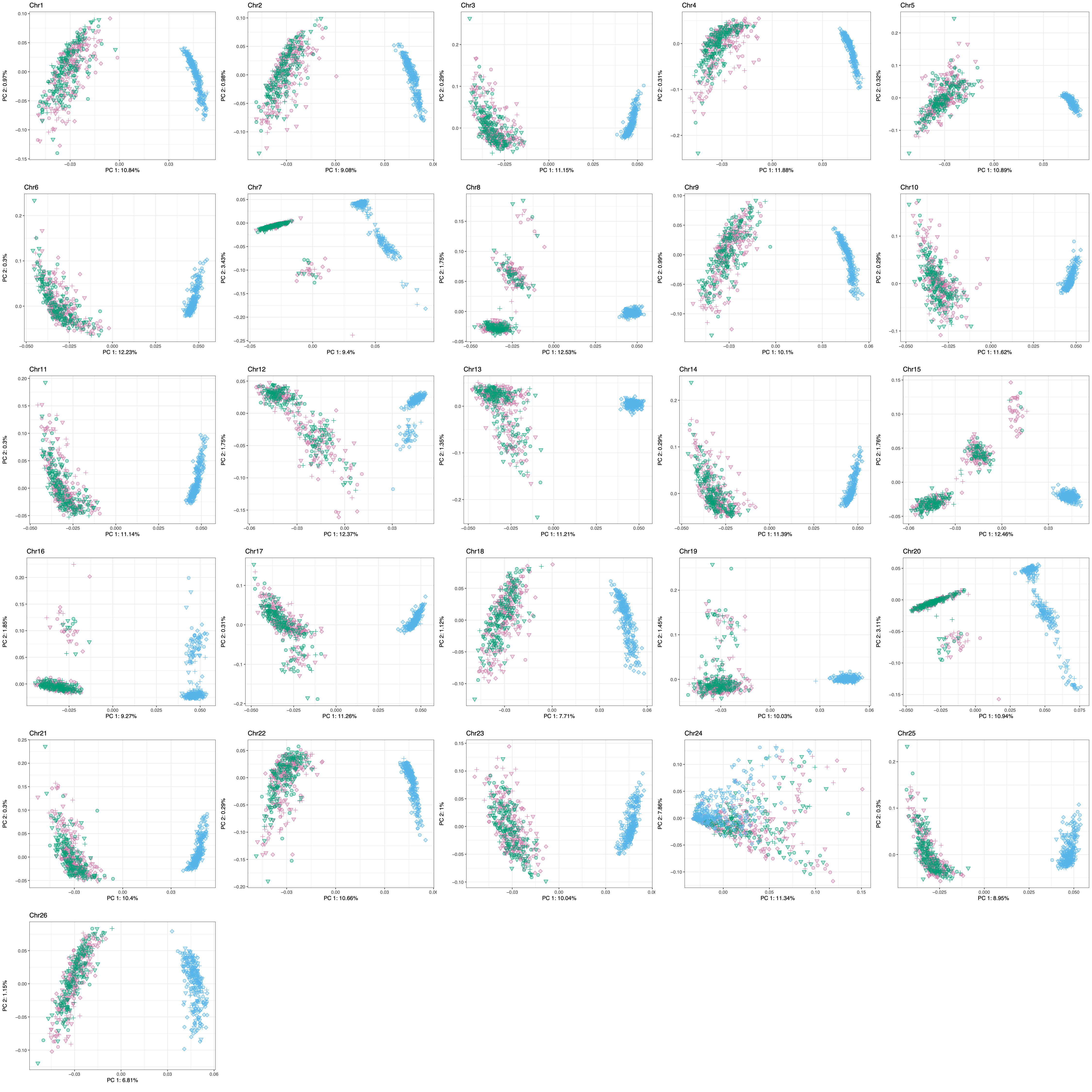

a

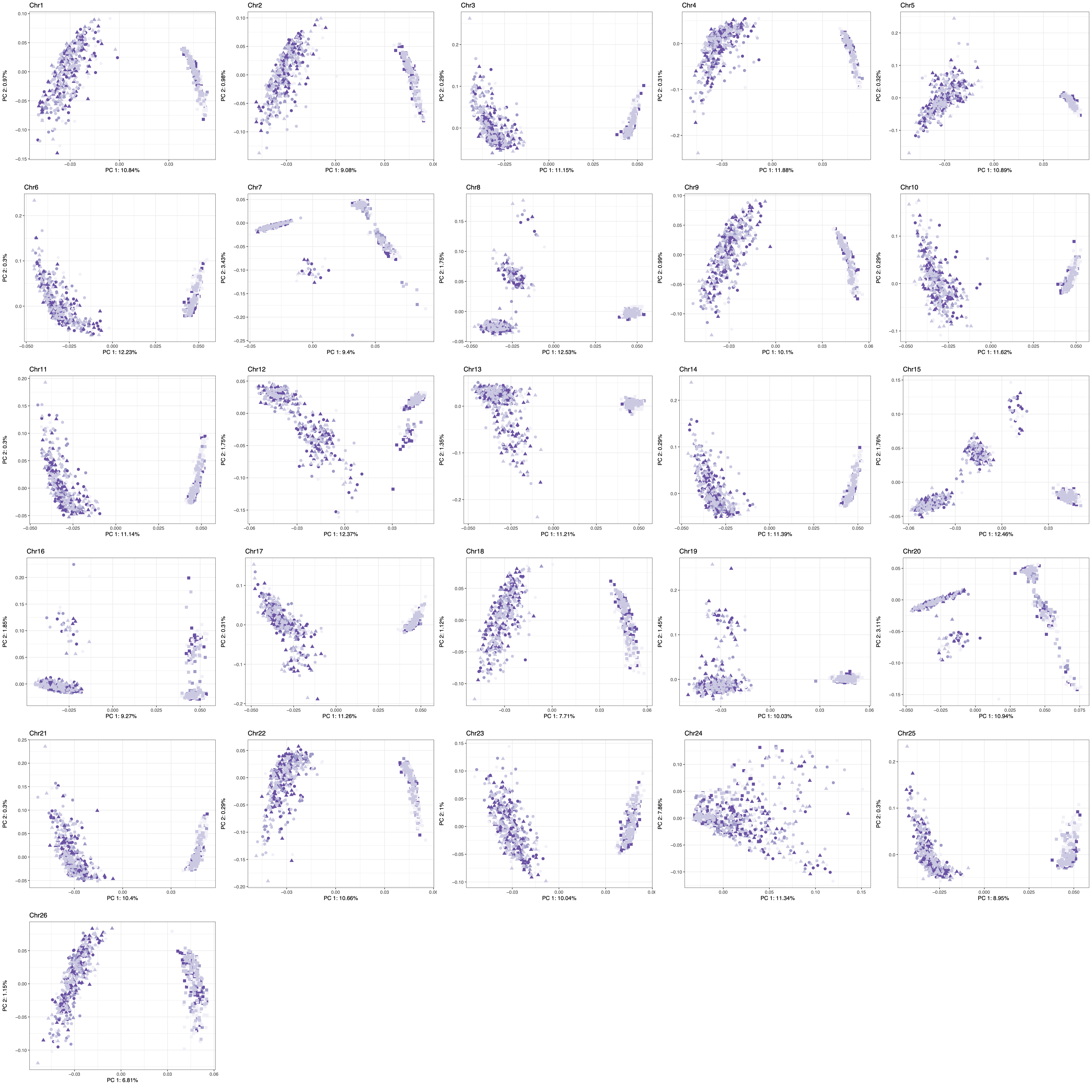

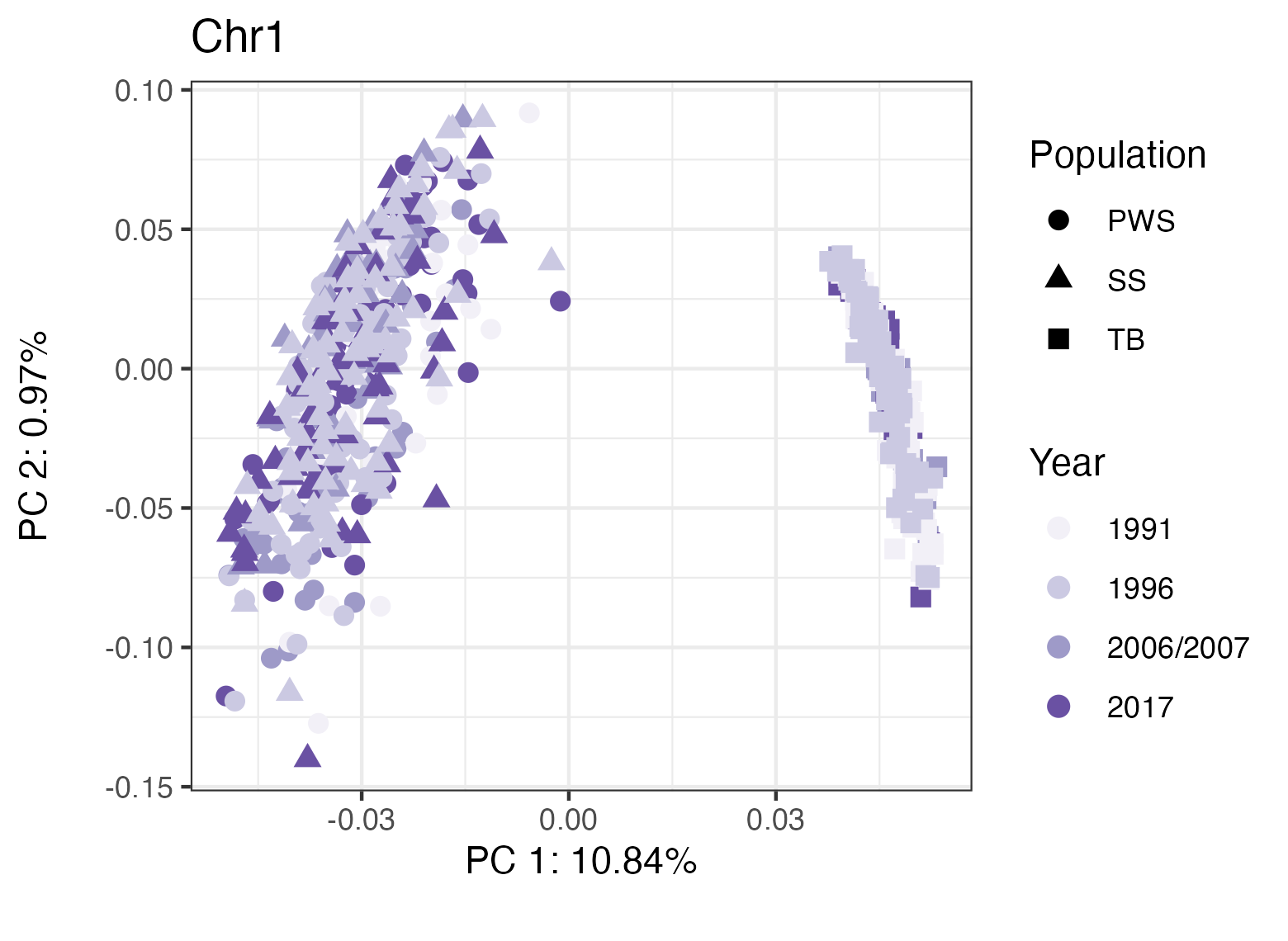

b

Fig. S1. PCA of each chromosome of Pacific herring populations. a) Colored by location/population, and b) colored by sampling year.

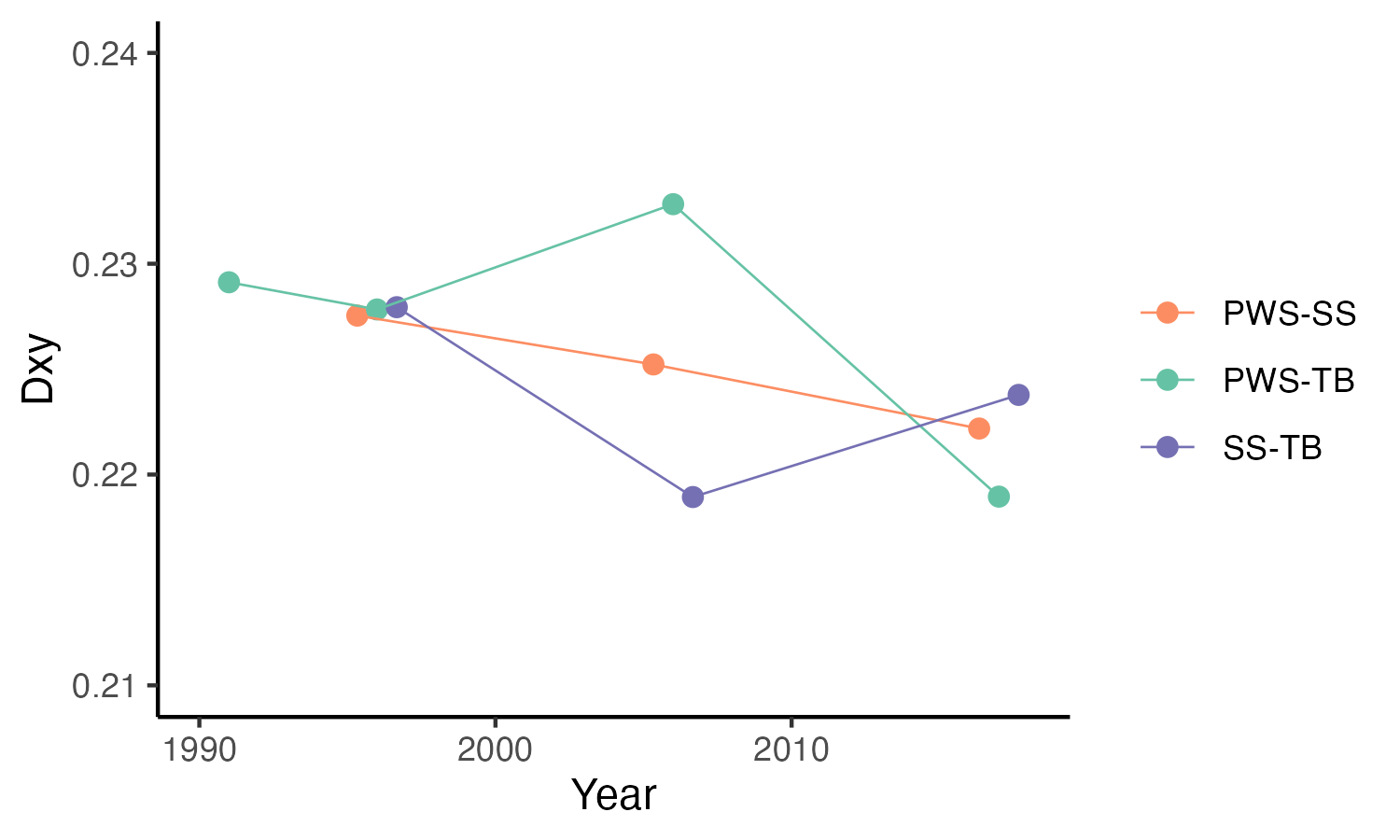

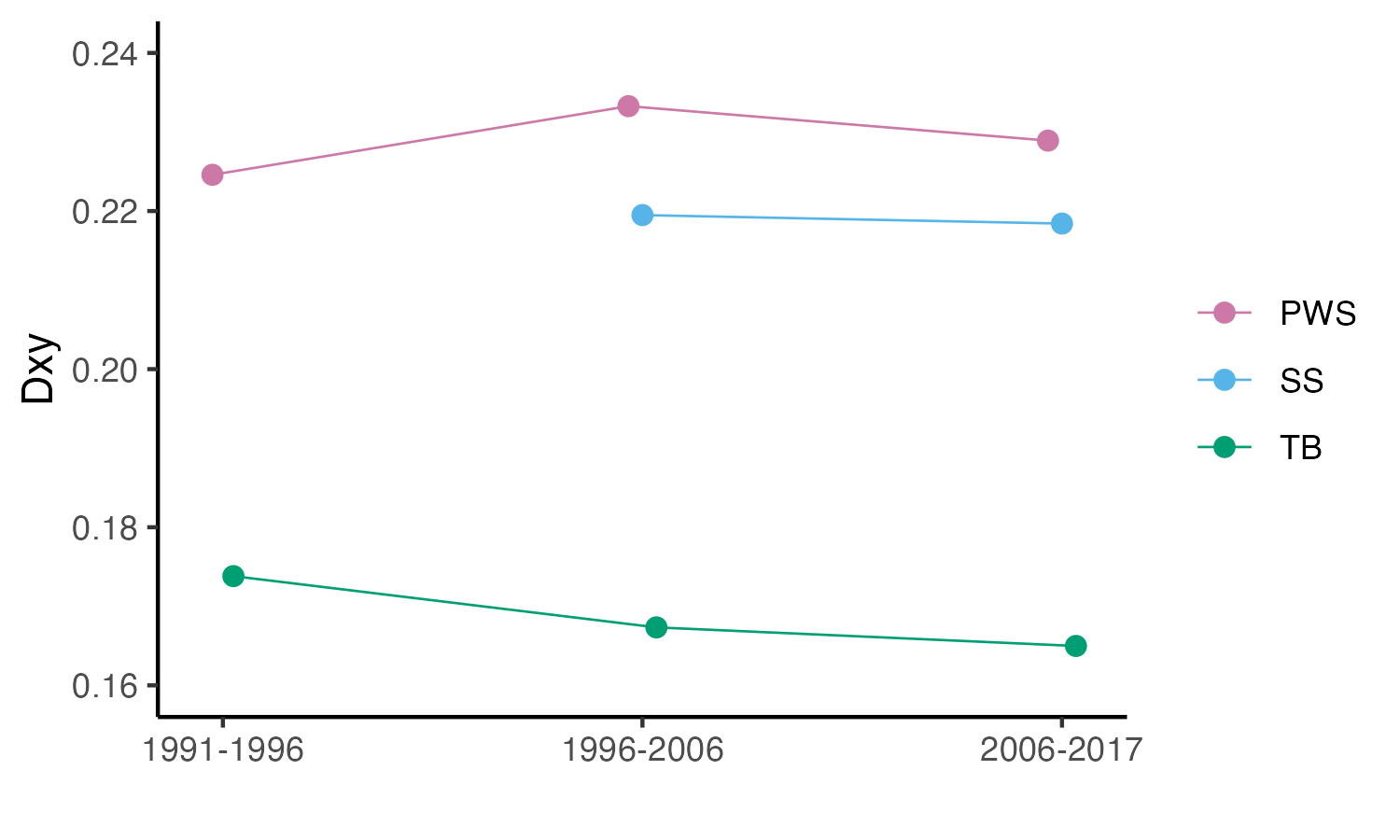

b

a

Fig. S2. D_XY_ estimated over time (a) within populations and (b) between populations.

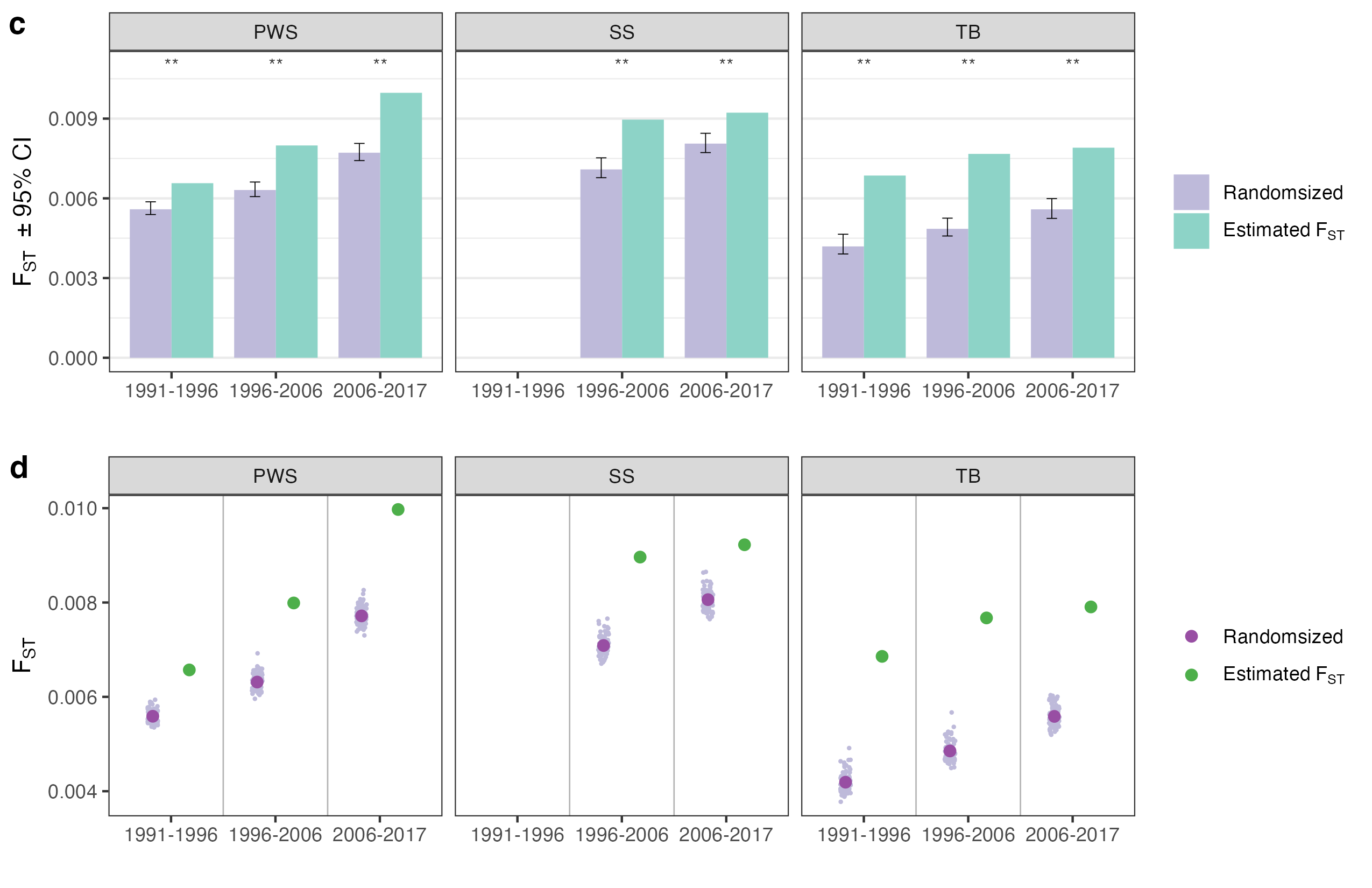

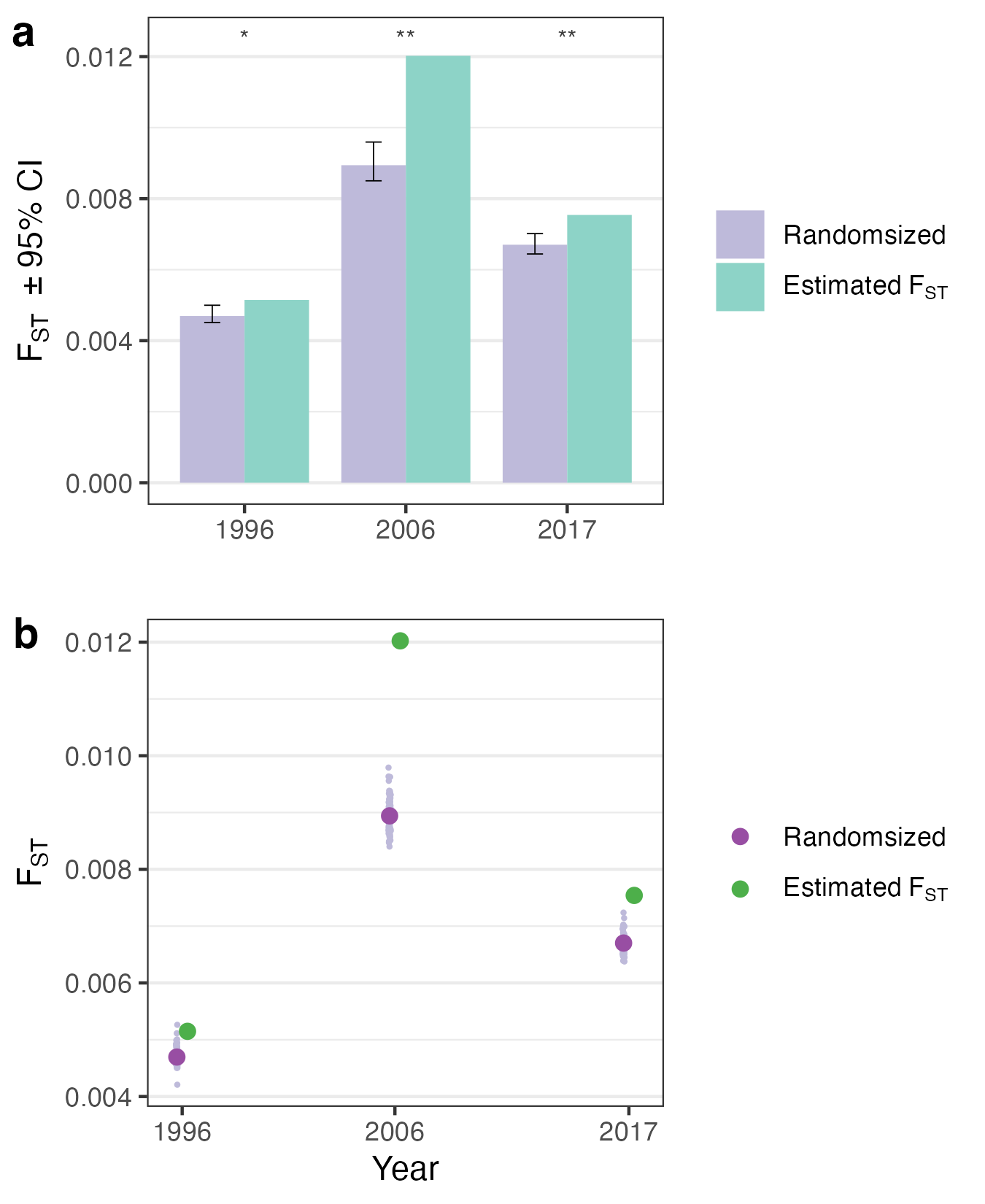

Fig. S3. F_ST_ values calculated from randomizing two groups vs. estimated values using ANGSD. a & b) Comparing F_ST_ between the PWS and SS populations at each time point. c & d) Comparing F_ST_ values over time points within each population. Asterisk symbols denote P-values (* = <0.05, ** = <0.01).

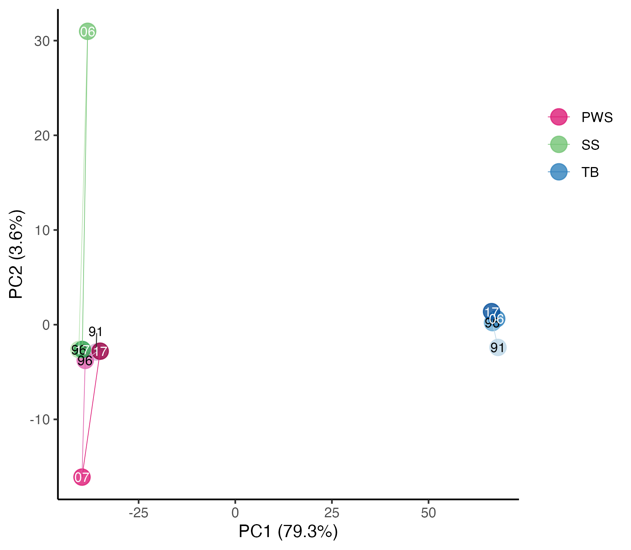

a

b

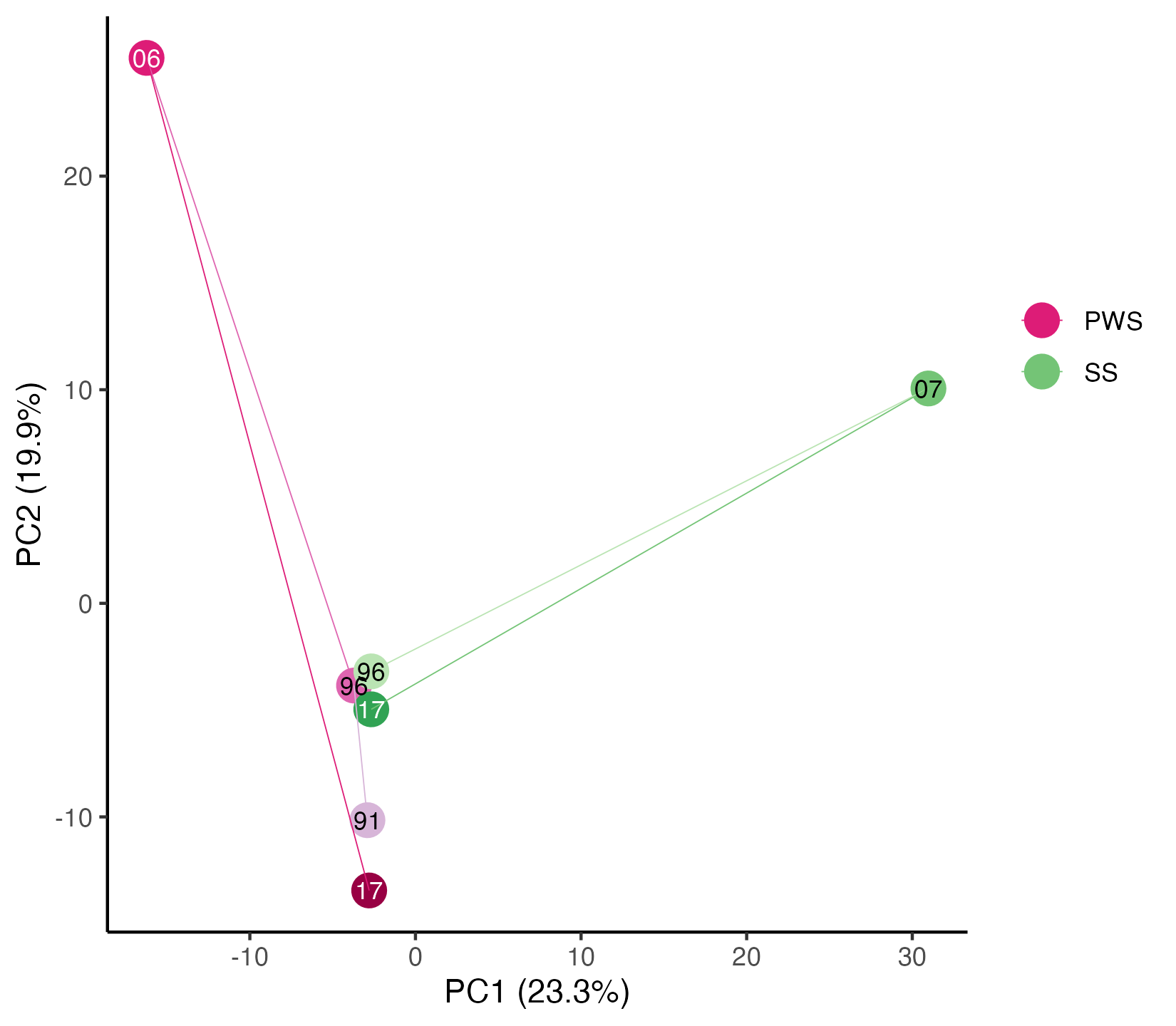

Fig. S4. PCA on the centered and standardized population frequencies at each time point. The connected series of points represent temporal shifts in population allele frequencies. a) A PCA of all three populations at all time points, showing a clear difference between the Bering Sea population (TB) and the Gulf of Alaska populations (PWS and SS). b) A PCA of PWS and SS populations only, highlighting large shifts in allele frequencies in 2006/07 in both populations.

Fig. S5. Results of PCAngsd selection scans for each Pacific herring population across all years. No loci were identified as significant (potentially selected) in any comparisons. We also tested for pairs of successive time points, but no significantly selected loci were identified (results not shown).

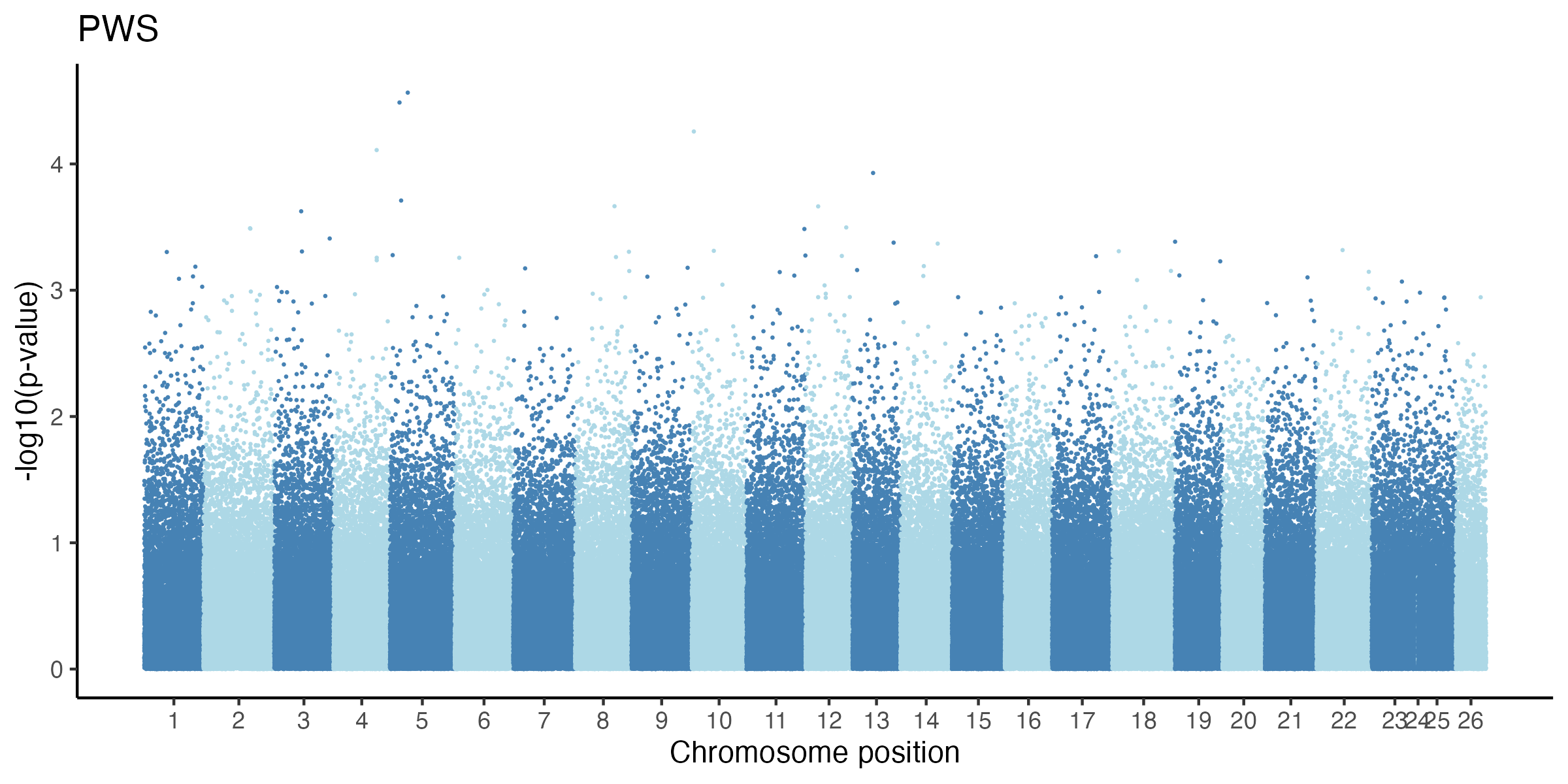

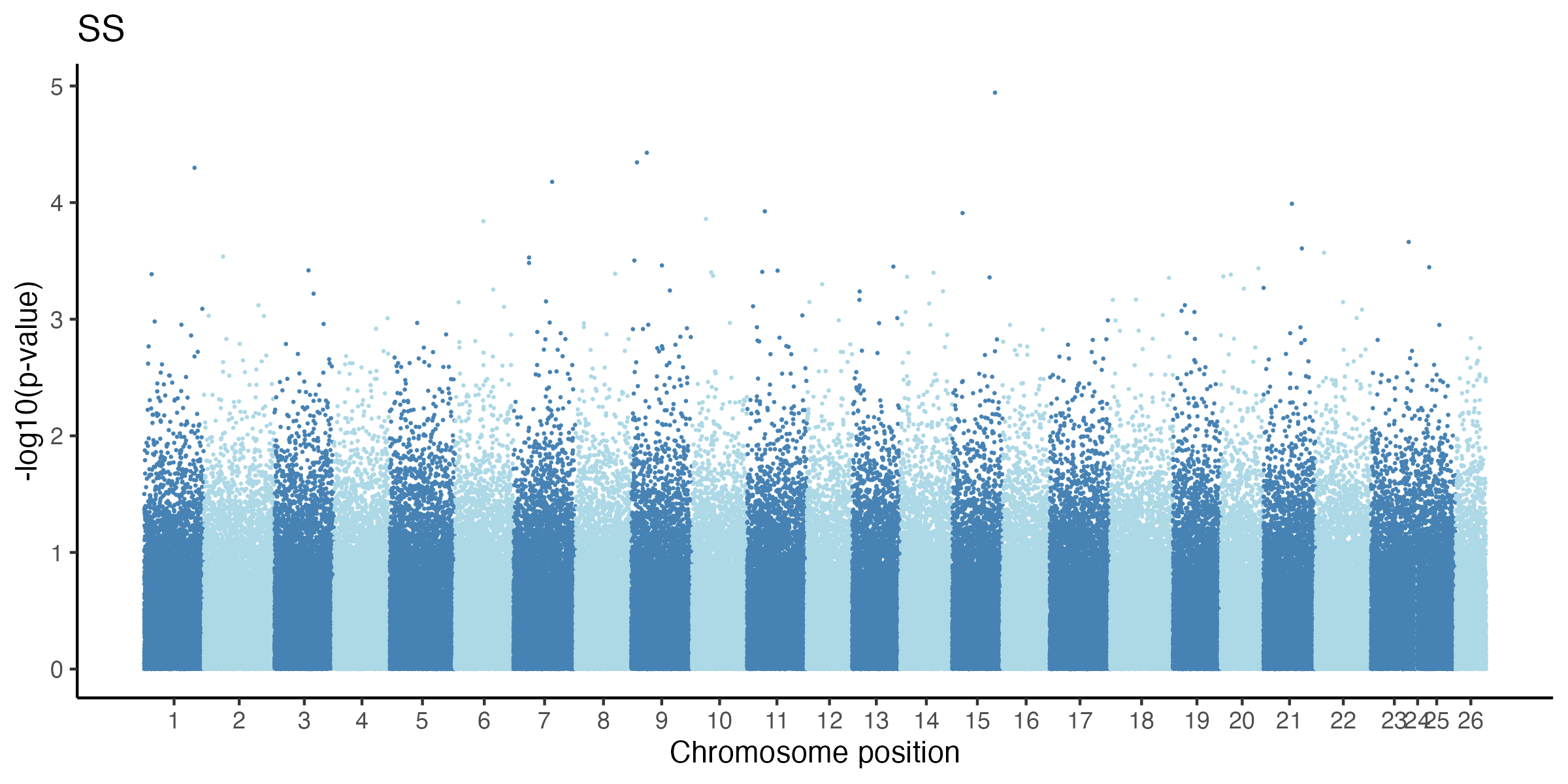

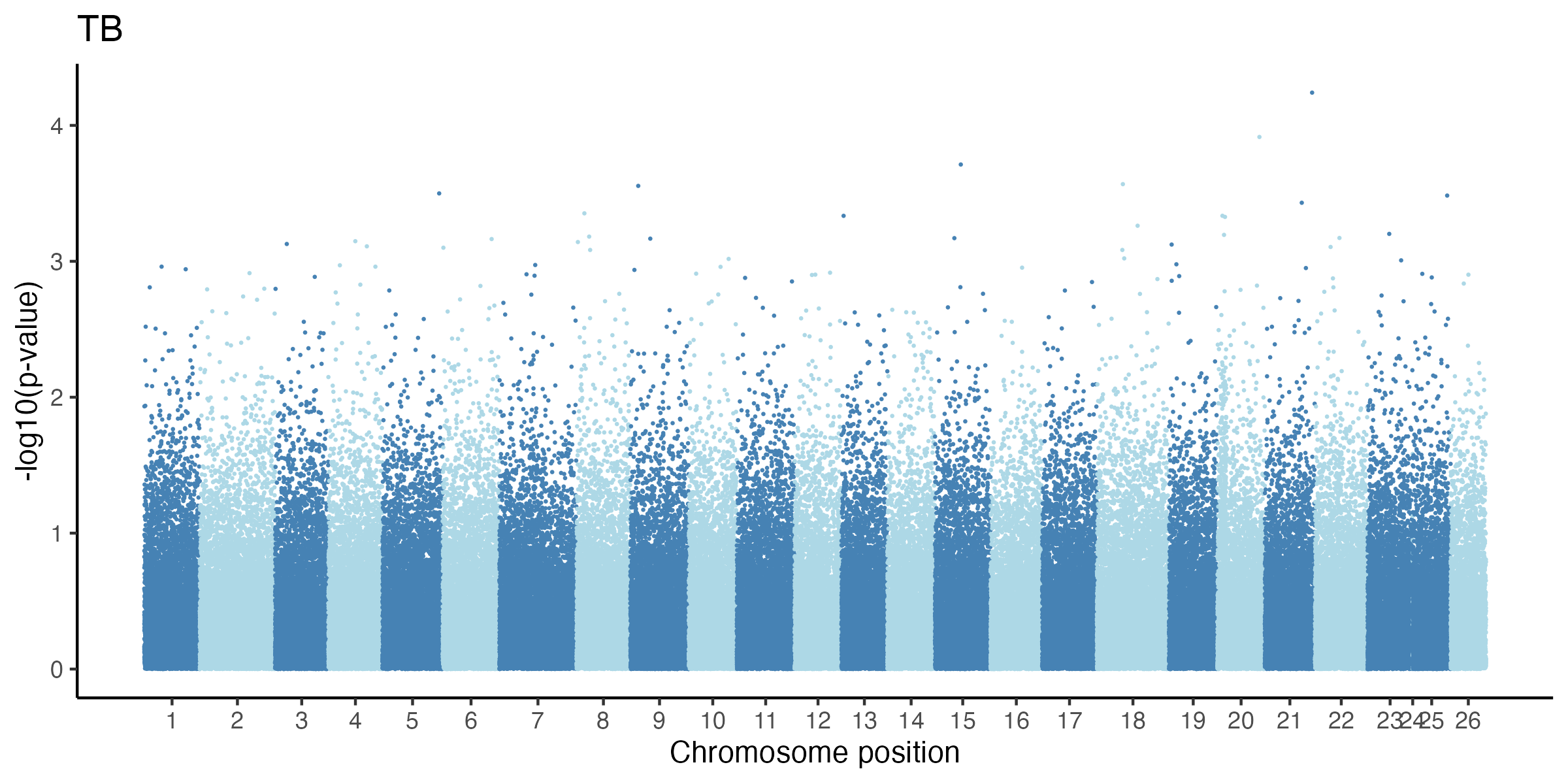

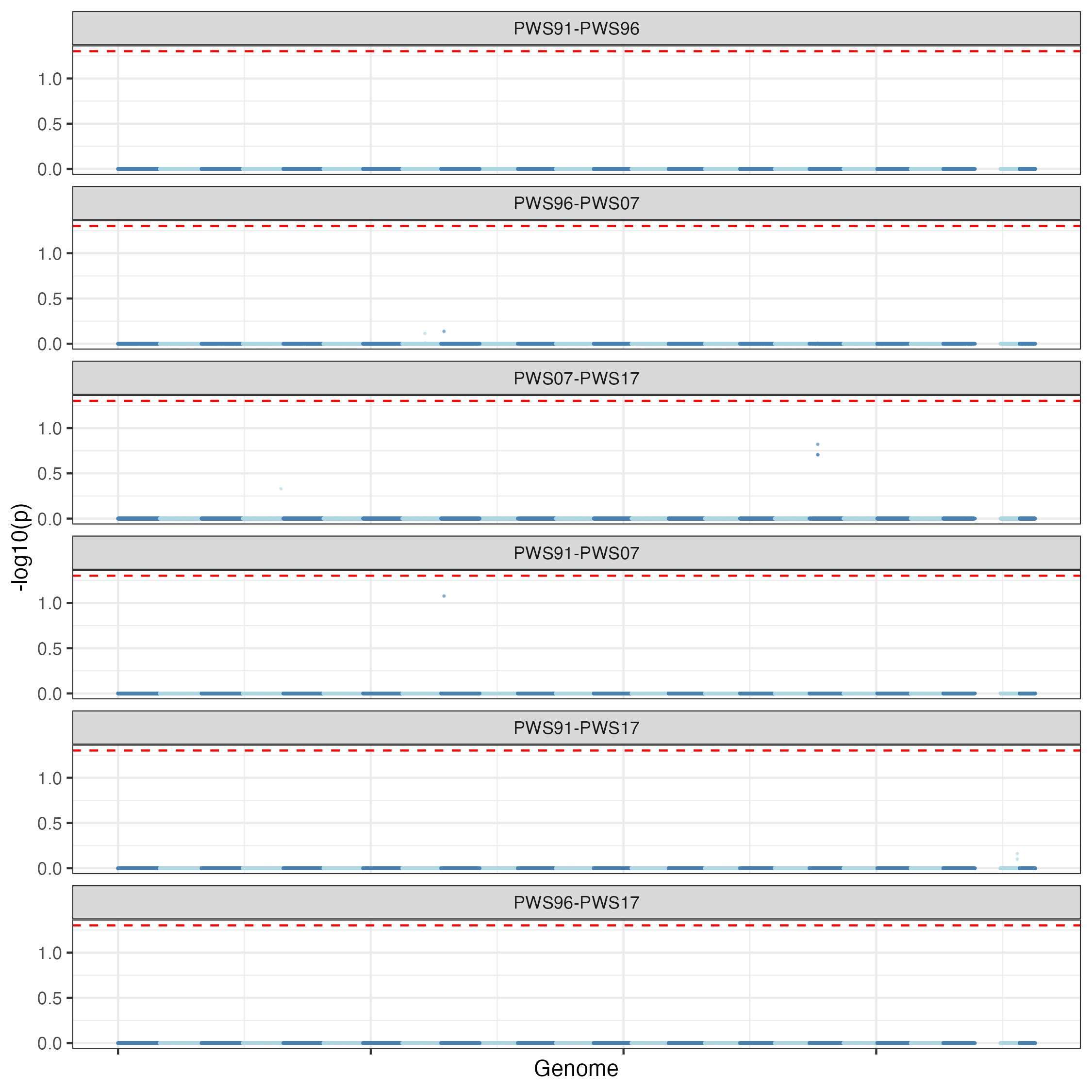

a

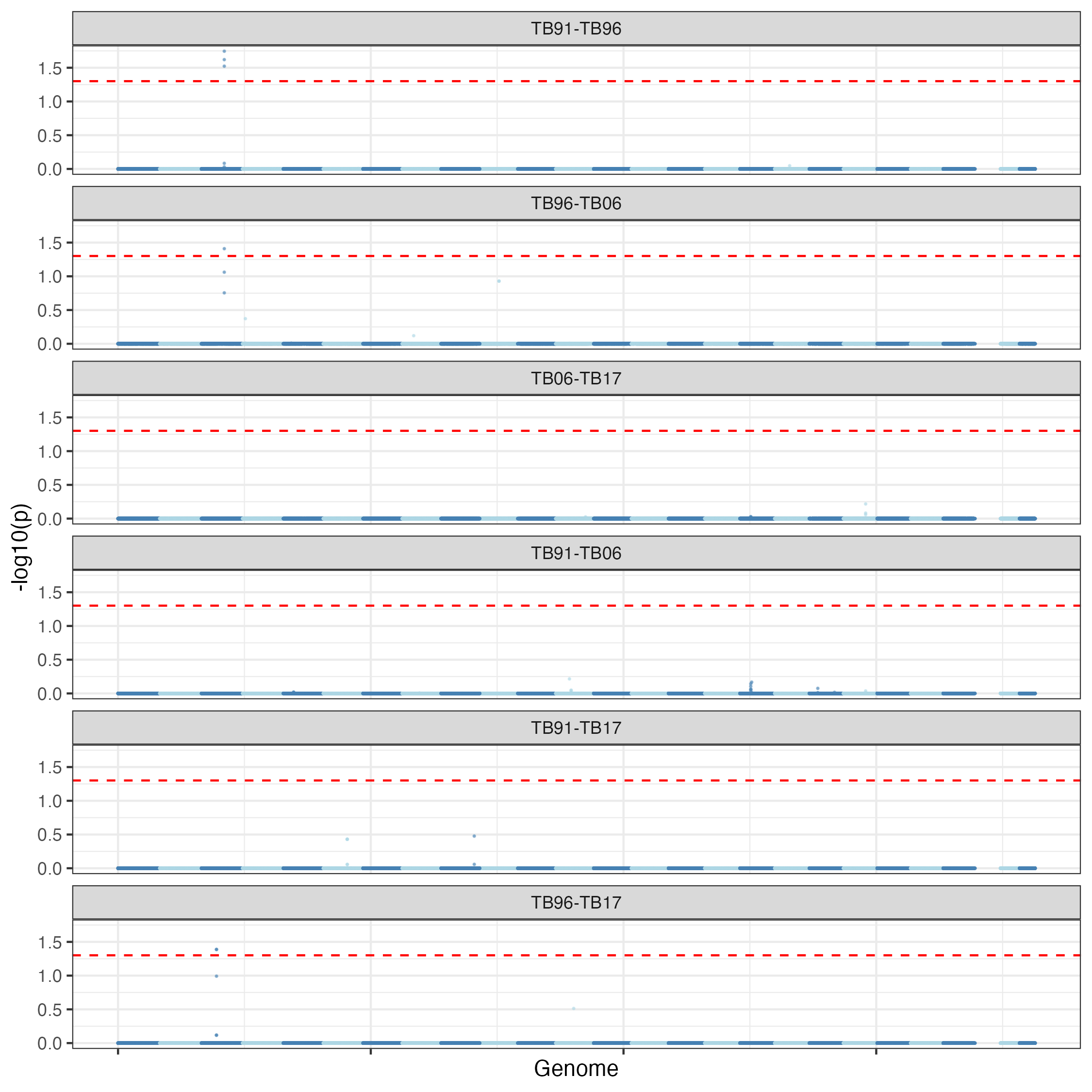

b

Fig. S6. Results of F_ST_ based genome scan using the method of Pinsky et al. (2021) performed by creating a null distribution of expected maximum F_ST_ for pairs of time points within a population. (a) The PWS population, (b) the TB population, and (c) the SS population. The red dot lines represent P-value <0.05. Further analysis showed that loci with relatively high P-values were all artifacts of windows with very few loci (<10).

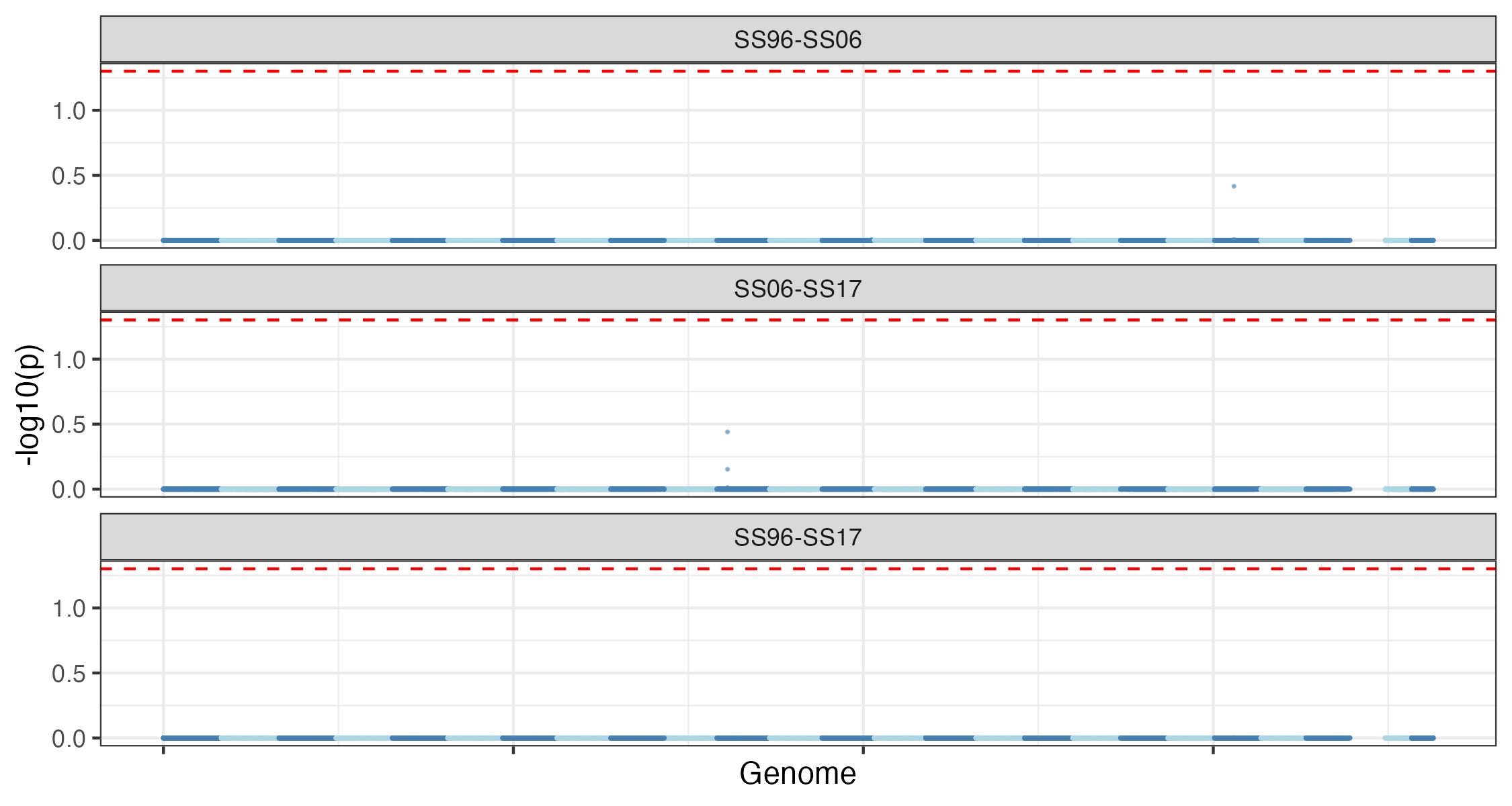

c

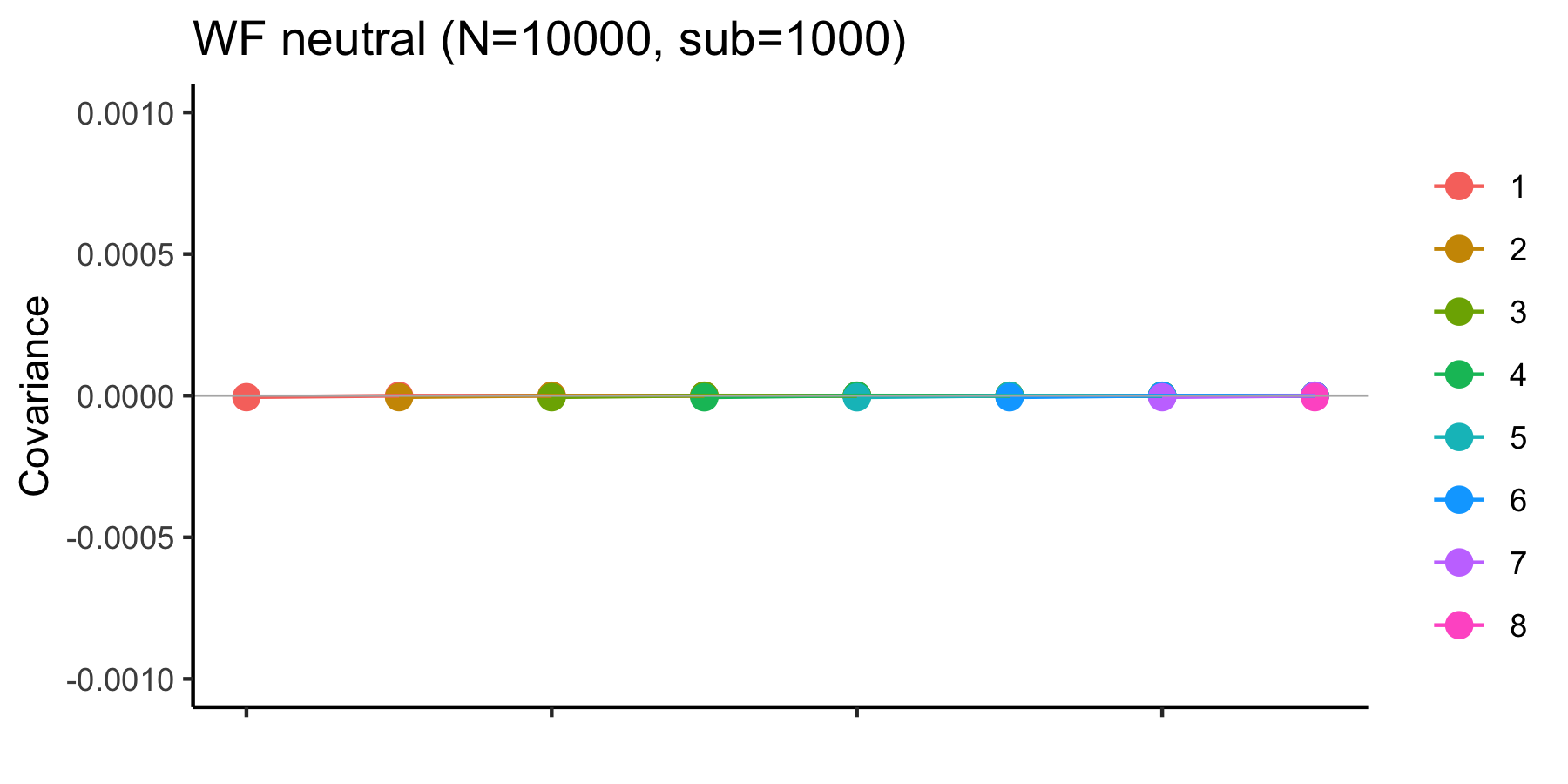

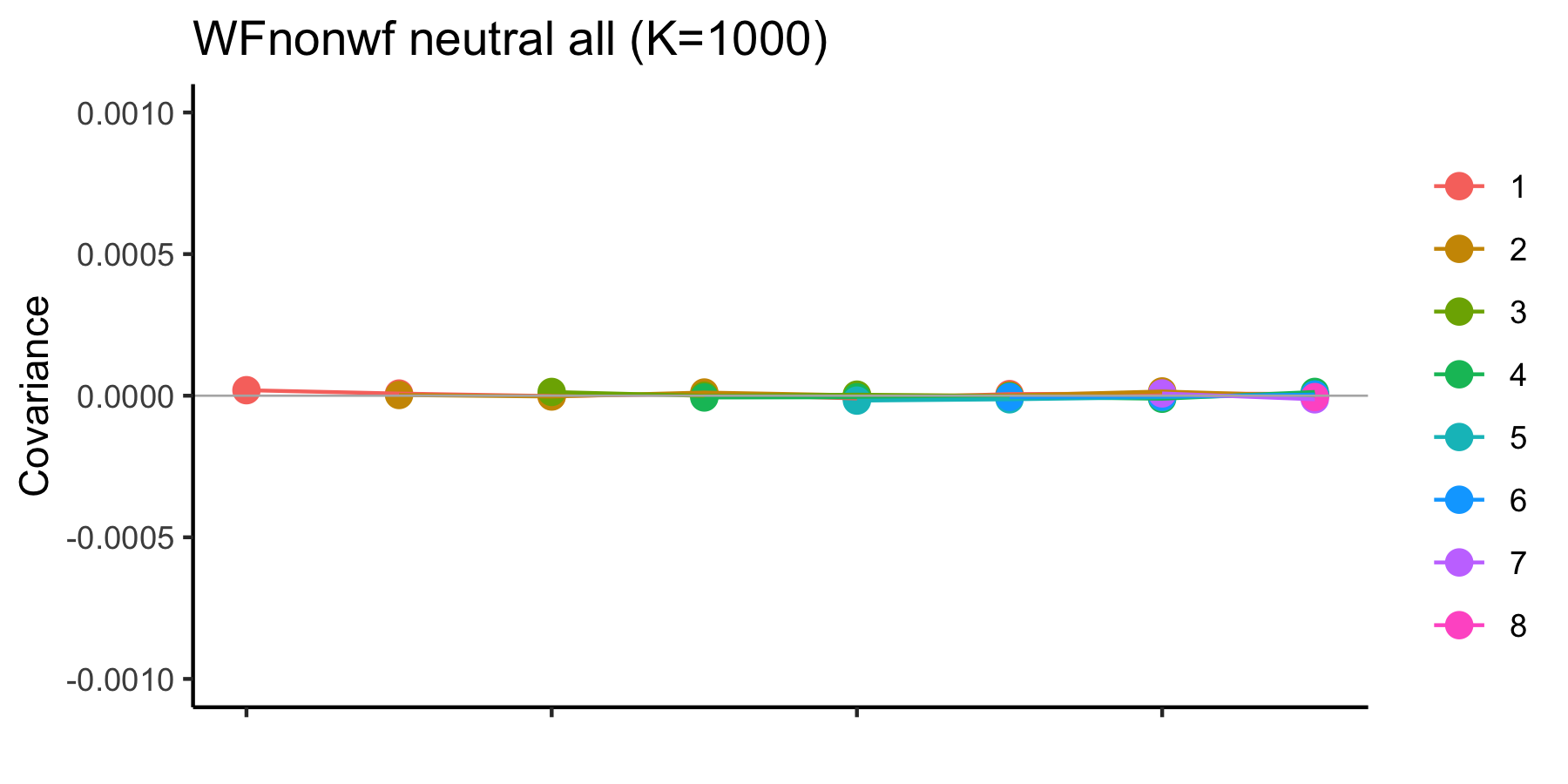

a

b

Fig. S7. Results from SLiM simulations comparing covariances of allele frequency changes using non-overlapping (a) and overlapping (b) generations in a neutral scenario over ten sampling times (i.e. eight covariances)*.* Simulations were run with the age first reproduction = 3 (at age 4) and sampling every 10 years. The results showed that the overlapping generations do not affect covariances. Hence *cvtk* can be applied to our data of Pacific herring with overlapping generations. a) Non-overlapping generation with N =10000 with S =1000, and b) overlapping generation with K=1000.

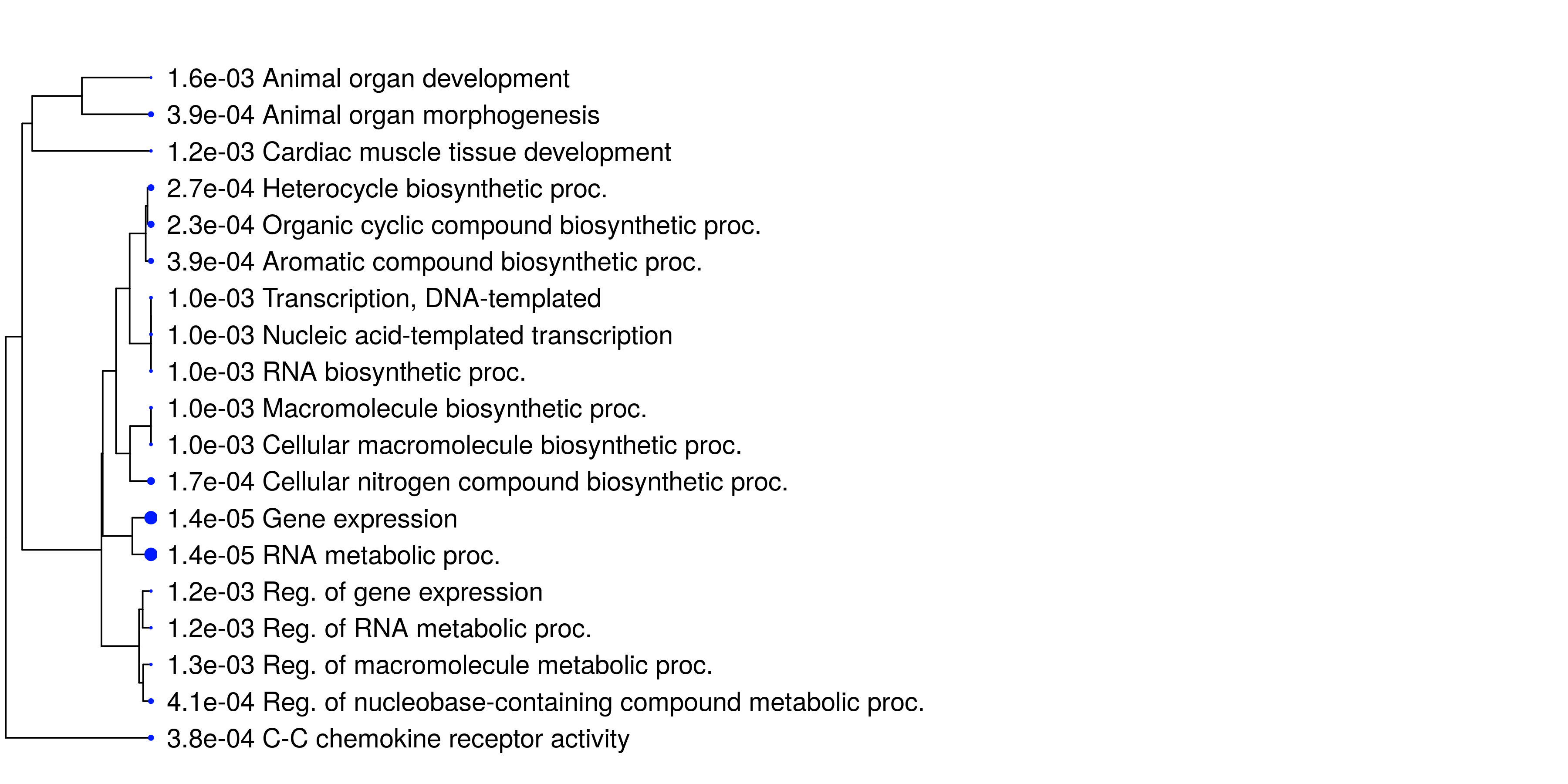

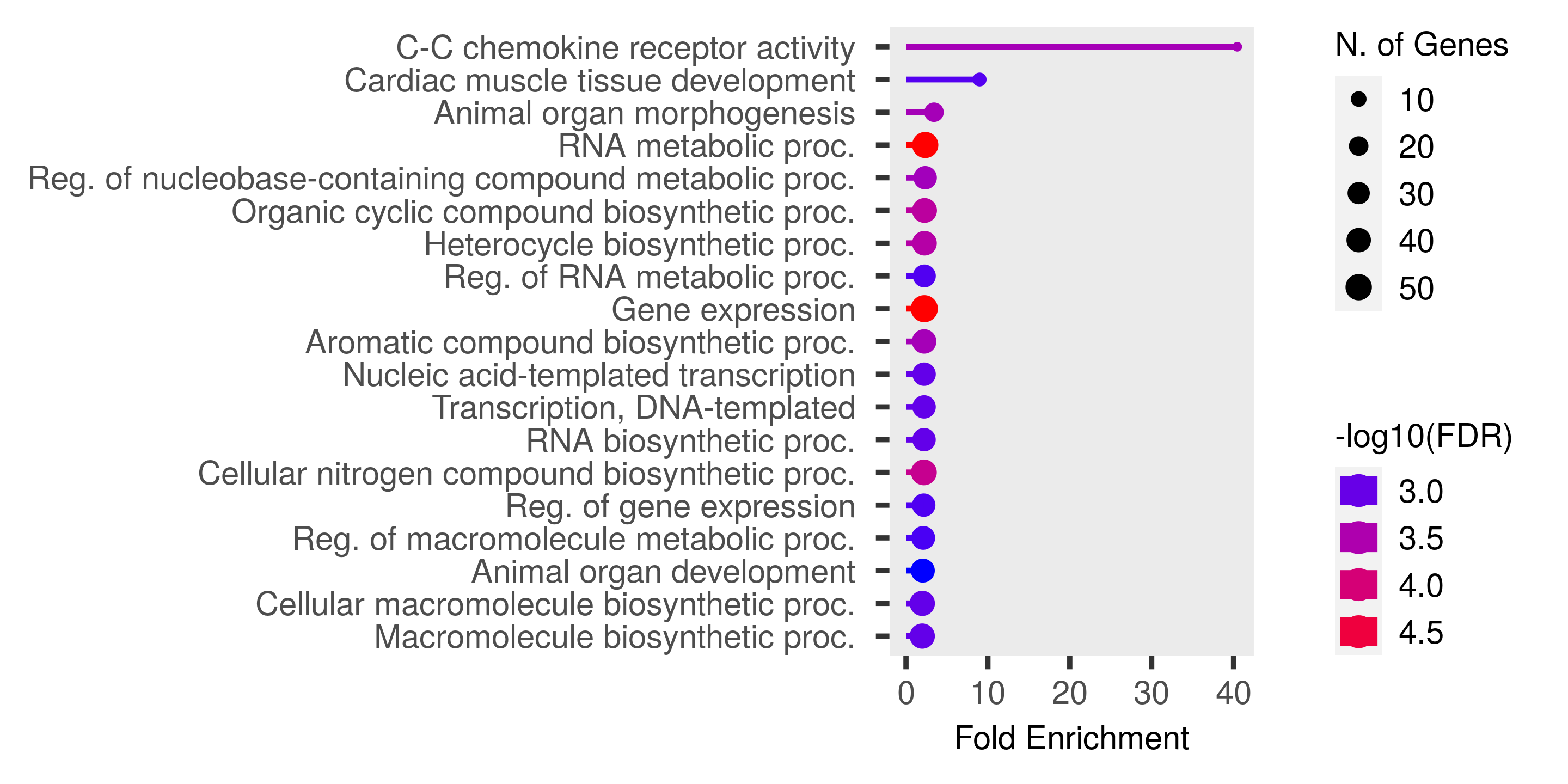

Fig. S8. Figures of Enriched GO terms generated by ShinyGO v.0.77 for the outlier regions (high covariances) in the PWS population for Period 1 (covariance between 1991-1996 and 1996-2007)
